# Inward transport of organelles drives outward migration of the spindle during *C. elegans* meiosis

**DOI:** 10.1101/2024.09.19.613972

**Authors:** Alma A. Peraza, Wenzhe Li, Aastha Lele, Denisa Lazureanu, Megan F. Hampton, Rebecca M. Do, Melissa C. Lafrades, Maria G. Barajas, Antonio A. Batres, Francis J. McNally

**Affiliations:** Department of Molecular and Cellular Biology University of California, Davis Davis, CA 95616

## Abstract

Cortical positioning of the meiotic spindle within an oocyte is required to expel chromosomes into polar bodies to generate a zygote with the correct number of chromosomes. In *C. elegans*, yolk granules and mitochondria are packed inward, away from the cortex while the spindle moves outward, both in a kinesin-dependent manner. The kinesin-dependent inward packing of yolk granules suggests the existence of microtubules with minus ends at the cortex and plus ends extending inward, making it unclear how kinesin moves the spindle outward. We hypothesized that inward packing of organelles might indirectly force the spindle outward by volume exclusion. To test this hypothesis, we generated a strain in which the only kinesin consists of motor domains with no cargo-binding tail optogenetically attached to mitochondria. This mitochondria-only kinesin packed mitochondria into a tight ball and efficiently moved the meiotic spindle to the cortex, supporting the volume exclusion hypothesis.

## Introduction

Oocyte meiosis is required to reduce the maternal genome copy number from 4 to 2 during meiosis I and from 2 to 1 during meiosis II. Accurate meiosis ensures a diploid copy number after fertilization by sperm whereas inaccurate oocyte meiosis leads to aneuploid or polyploid offspring, a common cause of pregnancy loss in humans^1^. Each meiotic division involves a meiotic spindle that is asymmetrically positioned with one spindle pole juxtaposed against the oocyte cortex. Asymmetrically positioned meiotic spindles induce extremely asymmetric cytokinesis that results in expulsion of chromosomes into tiny cells called polar bodies ^2^. Thus asymmetric positioning of oocyte meiotic spindles at the cortex is an essential part of animal development. In mouse, positioning of the meiotic spindle at the cortex requires formin 2-nucleated F-actin driving a mechanism that is still under active investigation ^3^. In contrast, positioning of the meiotic spindle in *C. elegans* requires microtubules ^4^ and the sequential action of kinesin 1 during prophase/prometaphase ^5^ and cytoplasmic dynein during anaphase ^6–8^. How two motor proteins that move in opposite directions on a microtubule can move the spindle in the same direction toward the oocyte cortex has remained unexplained.

*C. elegans* kinesin 1 heavy chain (UNC-116), kinesin light chain (KLC-1, 2), and kinesin cargo adapter (KCA-1) form a complex in vitro and have similar RNAi phenotypes ^5^, suggesting that they work together in vivo. In wild-type, the nucleus of the most mature prophase oocyte (-1 oocyte) often migrates to the cortex before nuclear envelope breakdown (NEBD) ^9^ then the spindle moves the remaining distance to the cortex ^4, 5^ during prometaphase as the oocyte ovulates through two sphincters, first into the spermatheca where fertilization occurs, then into the uterus where meiosis continues. We refer to metaphase I cells in the uterus as meiotic embryos, rather than oocytes, because they are already fertilized. Metaphase I spindles average 8 um in length with an elongated axial ratio of approximately 1.3 and adopt a mostly parallel orientation at the cortex. The anaphase promoting complex (APC) initiates sequential spindle shortening ^4^, dynein-dependent spindle rotation ^7^, anaphase I chromosome segregation and polar body cytokinesis. Dynein-dependent spindle rotation occurs when spindles have shortened to the shape of a sphere or slightly flattened sphere ^10, 11^. In meiotic embryos depleted of UNC-116 or KCA-1 by RNAi, elongated metaphase I spindles are several um away from the cortex, but move to the cortex in an APC-dependent ^5^ and dynein-dependent ^7, 8^ manner after the spindle shortens. After double depletions of UNC-116 and dynein, spindles shorten far from the cortex ^7^. In contrast, in *unc-116(f130)* meiotic embryos, metaphase spindles are further from the cortex than in RNAi embryos, and spindle shortening often proceeds in the middle of the embryo ^10, 12^. Thus it has been unclear whether *unc-116(f130)* causes a neomorphic spindle centering phenotype or if RNAi can lead to incomplete depletion.

In *C. elegans* prophase -1 oocytes, yolk granules and mitochondria are evenly dispersed. In metaphase I meiotic embryos, yolk granules ^13^ and mitochondria ^14^ are packed inward, away from the cortex and both organelles begin circular cytoplasmic streaming. Inward packing of yolk granules and mitochondria, as well as cytoplasmic streaming are dependent on UNC-116 and KCA-1 ^13^. Kinesin-dependent inward packing suggests the existence of microtubules with minus ends at the cortex and plus ends extending inward. However, it has been unclear if outward spindle translocation, inward packing of yolk granules and mitochondria, and cytoplasmic streaming are due to kinesin binding to different cargos. The C-terminus of kinesin heavy chain can bind many different cargos either directly or through a light chain ^15^. For example, in *C. elegans* axons, UNC-116 is targeted to mitochondria through RIC-7 ^16^ or the redundant activities of MIRO and metaxin ^17^, whereas in hyp7 cells, UNC-116 is targeted to the nuclear envelope by UNC-83 ^18^. The situation in meiotic embryos is particularly enigmatic because the spindle moves outward, in the opposite direction of the inward packing yolk granules and mitochondria. There might be two populations of microtubules with opposite orientation or one movement might be indirectly caused by the motor-driven movement in the opposite direction. Here we test the hypothesis that motor-driven inward packing of organelles pushes the spindle outward by volume exclusion.

## Results

### A kinesin-1 germline null allele leads to a centered meiotic spindle during metaphase I

Because *unc-116(RNAi)* previously resulted in a less severe spindle positioning defect ^5^ than the *unc-116(f130)* ts allele ^12, 19^, we first sought to establish the germline null phenotype for *unc-116* (kinesin 1 heavy chain). The *unc-116(gk5722)* allele ^20^ is a larval lethal CRISPR deletion of the 5’ two thirds of the *unc-116* locus and thus should be a null allele. We generated strains in which this allele is complemented in somatic cells by an integrated repetitive array of *unc-116::GFP (duIs1)* that is silenced in the germline to allow observation of meiotic embryos that lack UNC-116. When control meiosis was monitored by time-lapse in utero imaging, 10/10 metaphase I spindles (indicated by their elongated shape and timing after ovulation into the uterus) were located at the cortex and mitochondria were packed inward away from the cortex (Fig. 1A). In contrast, 13/13 *unc-116(gk5722) duIs1* metaphase I spindles were centered in the embryo (Fig. 1B, 1C) much further from the cortex than previously reported for *unc-116(RNAi)* ^5^. Anaphase promoting complex (APC) activation initiates sequential spindle shortening, dynein-dependent spindle rotation (Fig. 1A), then anaphase chromosome separation. In UNC-116-depleted embryos, this dynein-dependent pathway can mediate late translocation to the cortex ^5, 7^. 6/13 shortened anaphase *unc-116(gk5722) duIs1* spindles moved to the cortex late (Fig. 1B) whereas 7/13 remained centered resulting in pentaploid embryos (4 maternal + 1 paternal genome)(Fig. 1C). 18/18 *unc-116(f130ts)* metaphase I spindles incubated at nonpermissive temperature were also centered as previously described ^12, 19^(Fig. 1D, 1E, 1F) and a fraction of these spindles underwent late translocation (Fig. 1E) whereas a fraction remained centered (Fig. 1F) resulting in pentaploid embryos. Previous work ^7, 8^ indicated that late translocation after UNC-116 depletion is due to a separate dynein-dependent mechanism rather than residual kinesin-1 activity. We therefore refer to *unc-116(gk5722) duIs1* as *unc-116(g-null)* for germline null.

**Figure 1.**
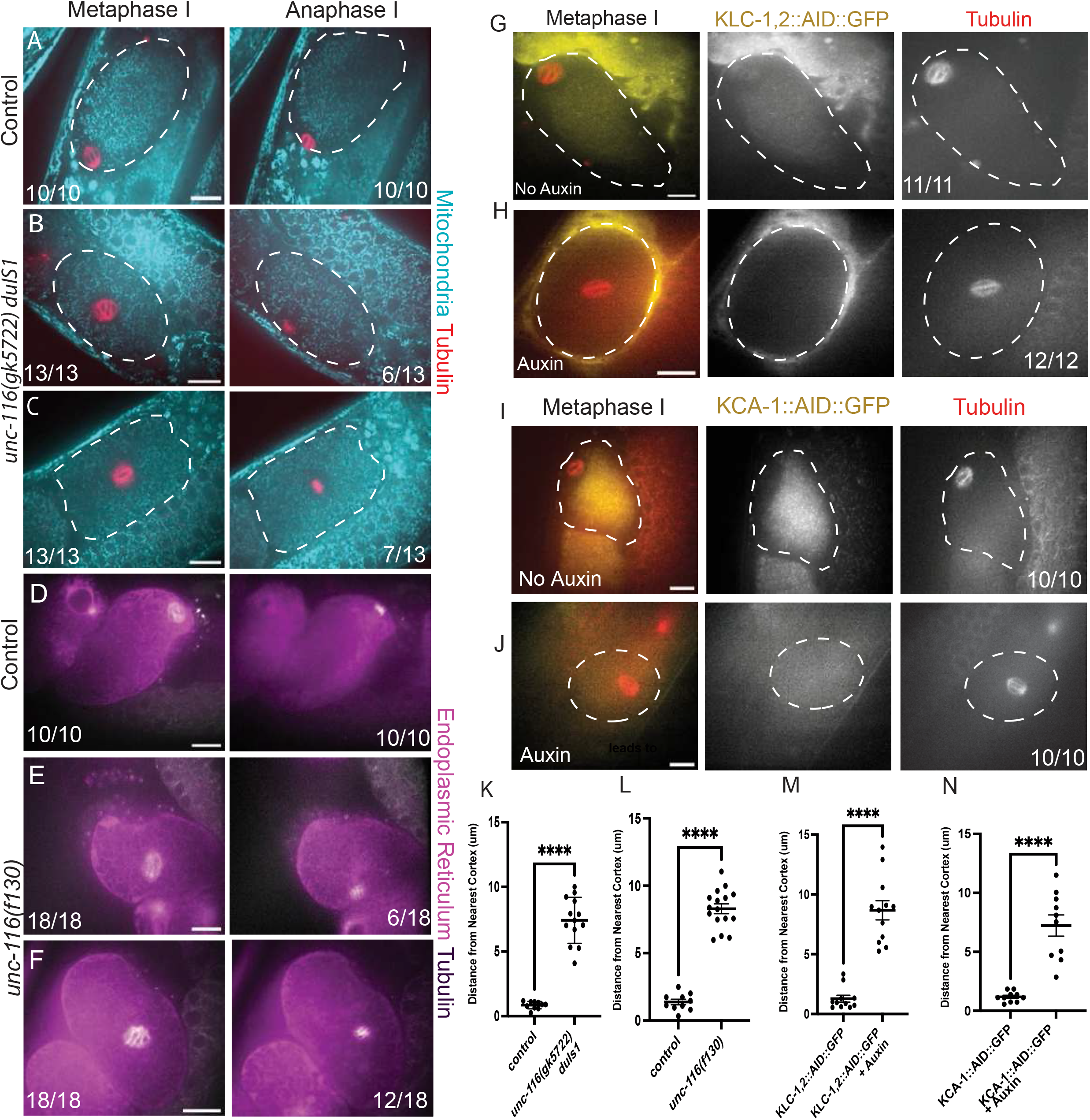
***unc-116(gk5722) duIs1*, *unc-116(f130 ts)*, or auxin-induced degradation of KLC-1,2 or KCA-1 lead to a centered metaphase I spindle.** Single plane time-lapse in utero imaging of anesthetized *C. elegans*. A. Control meiotic embryo expressing COX-4::GFP labeled mitochondria (cyan) and mKate::tubulin (red). Mitochondria are packed inward and the spindle is located at the cortex (n=10/10). B and C. *unc-116(gk5722) duIs1* germline null gives rise to a centered spindle during metaphase I but has two distinct spindle localization phenotypes during anaphase I. 6/13 spindles underwent late translocation (B) and 7/13 remained centered (C). D. Control meiotic embryo expressing SP12::GFP labeled ER (magenta) and mKate::tubulin (grayscale). Metaphase I spindle is at the cortex (n= 10/10). E and F. *unc-116(f130ts)* has a centered spindle during metaphase I. During anaphase I, 6/18 spindles underwent late translocation (E) and 12/18 remained centered. G. Metaphase I spindles are at the cortex in KLC-1,2::AID::GFP embryos without auxin (n=11/11). H. Auxin treated KLC-1,2::AID::GFP metaphase I embryos have centered spindles (n=12/12). I. Metaphase I spindles are at the cortex of KCA-1::AID::GFP embryos without auxin (n=9/9). J. Auxin treated KCA-1::AID::GFP metaphase I embryos have centered spindles (n=10/10). K-N. Distance from the edge of the metaphase I spindle to the nearest point on the cortex. Bar = 10 μm. **** indicate p < .0001 Student’s t test.

In a previous study, either double depletion of both *C. elegans* kinesin-1 light chains or single depletion of KCA-1 by RNAi resulted in metaphase I spindles that were several um from the cortex but not quite centered ^5^. To validate these RNAi results, we used CRISPR to tag the endogenous *klc-1*, *klc-2* and *kca-1* genes with auxin induced degrons (AID) and GFP. Auxin addition led to the reduction of KLC-1,2::AID::GFP fluorescence and resulted in 12/12 centered metaphase I spindles (Figure 1G, 1H).

Similarly, auxin addition led to reduced KCA-1::AID::GFP fluorescence and 10/10 centered metaphase I spindles (Figure 1I, 1J). Quantification of metaphase I spindle distance to the nearest cortex is shown in Fig. 1K-1N. These results indicate that a complete loss of UNC-116, KLC-1 + KLC-2, or KCA-1 results in a centered metaphase I spindle.

### KCA-1 and KLC-1 are localized on yolk granules which pack inward at the same time that the spindle moves outward

A previous study found that yolk granules are evenly distributed in -1 oocytes before nuclear envelope breakdown and packed inward during metaphase I after ovulation. In addition, UNC-116, KCA-1 and KLC-1 were observed only in a gradient in the cytoplasm of metaphase I embryos with higher concentration in the middle of the embryo (McNally 2010). If inward yolk granule movement causes outward spindle movement, it should occur simultaneously with spindle movement and kinesin should be localized on organelles during outward spindle movement. To test these possibilities, oocytes were monitored by time-lapse imaging from before nuclear envelope breakdown in the gonad, during ovulation through the spermatheca, through metaphase I in the uterus. In 10/10 control worms, yolk granules labelled with endogenously tagged VIT-2::GFP were evenly distributed throughout the oocyte before nuclear envelope breakdown (Figure 2A; -10:58). During NEBD, indicated by assembly of mKate::tubulin into a spindle within the nuclear volume, the yolk granules began packing inward away from the mCherry::PH-labelled plasma membrane (Fig. 2A; 12:58). Inward packing of yolk granules and outward movement of the spindle occurred simultaneously while the zygote was in the spermatheca (Fig. 2A; 13:41) and the metaphase I spindle was fully positioned at the cortex by the time the embryo exited the spermatheca into the uterus (Fig. 2A; 18:34). In 12/12 *unc-116(f130)* ts mutants, the yolk granules remained evenly dispersed and the spindle remained in the center of the embryo (Figure 2B; 12:05). In 19/19 worms with endogenously tagged KCA-1::AID::GFP and endogenously tagged VIT-2::HALO, KCA-1 appeared to co-localize with VIT-2-labelled yolk granules that were evenly dispersed before NEBD (Fig. 2C, 2D; -8 min, Video S1). In 11/11 ovulating oocytes, KCA-1 on yolk granules packed inward during spindle assembly, ovulation, and spindle migration but dispersed from the yolk granules after yolk granule packing and spindle migration (Fig. 2C, 2D; compare 2C’ vs 2C’’, Video S1). Endogenously tagged KLC-1::AID::GFP also co-localized with VIT-2::HALO before and during yolk granule packing and spindle migration (Fig. 2E, 2F; -16:50; Fig. S1C, Video S2) and also dispersed from yolk granules after ovulation when the spindle was already at the cortex (Fig. 2E, 2F; compare 2E’ vs 2E’’, Video S2). The high density of yolk granules made definitive co-localization of KCA-1 with VIT-2 challenging (Fig. 2H, 2I). To more rigorously test the idea that KCA-1 is on the surface of yolk granules, we depleted RAB-7 by RNAi, which causes yolk granules to become much larger ^21, 22^. KCA-1 appeared as a ring surrounding the enlarged VIT-2-labelled yolk granules in single plane optical sections of *rab-7(RNAi)* oocytes before NEBD (Fig. 2J, 2K). KCA-1::AID:GFP and KLC-1::AID::GFP fluorescence were clearly anti-correlated with a mitochondrial marker (Fig. S1), further supporting the conclusion that the greatest enrichment of kinesin-1 is on the surface of yolk granules, instead of mitochondria during migration of the spindle to the cortex.

**Figure 2.**
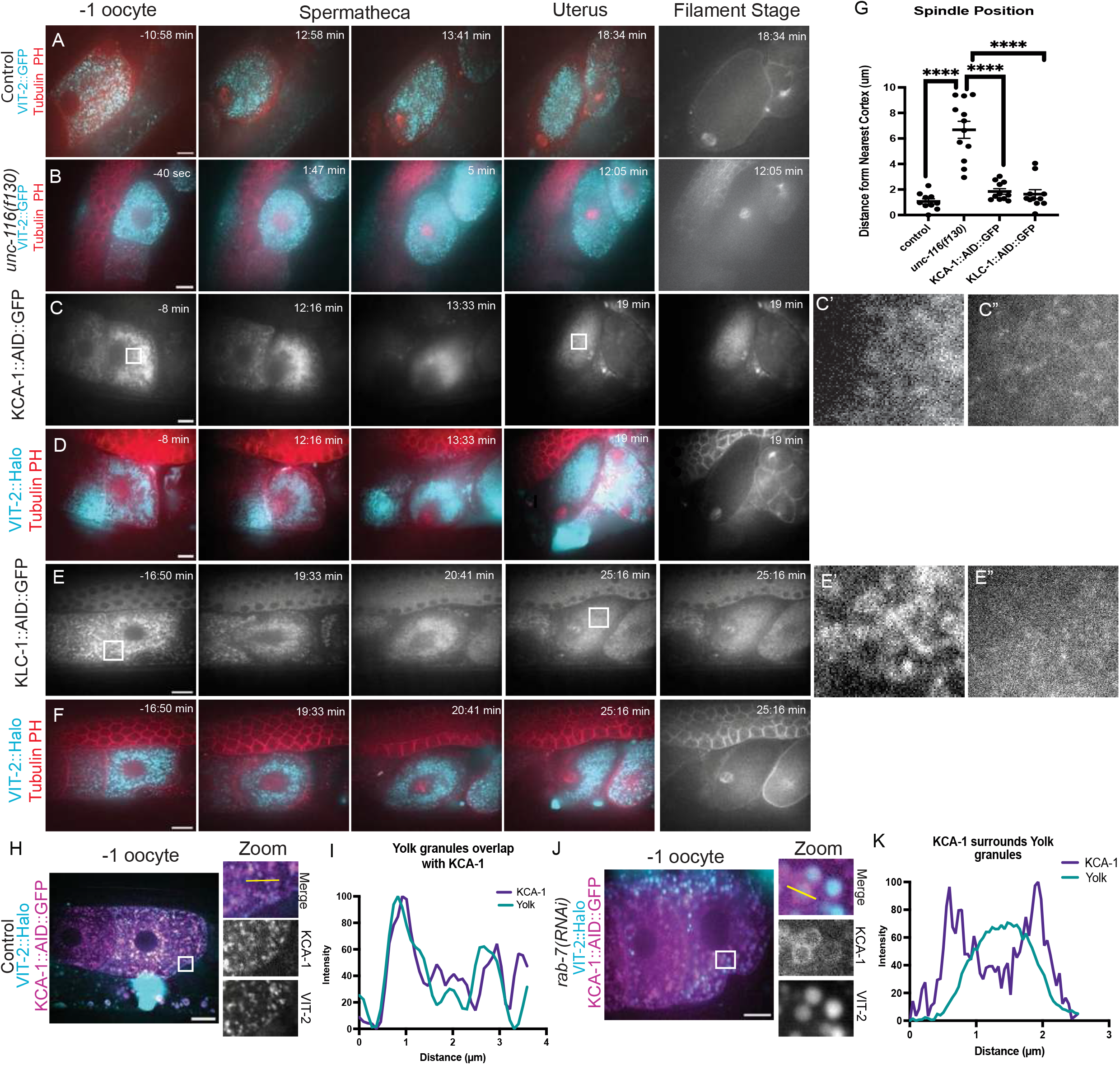
Yolk granules pack inward as the spindle moves outward and, KCA-1 and KLC-1 co-localize with yolk granules. Single plane time-lapse in utero imaging of anesthetized *C. elegans*. A. Control oocyte with VIT-2:GFP labeled yolk granules (cyan), mKate::tubulin (red) and PH (plasma membrane-red). Yolk granules are dispersed in the -1 oocyte, then pack inward at the same time the spindle moves outward. B. *unc-116(f130)ts* oocyte with yolk granules (cyan) that do not pack inward and a spindle (red) that remains centered. C and D. Before ovulation (-8 min), KCA-1::AID::GFP (grayscale) is on vesicles that co-localize with VIT-2::Halo (cyan). During ovulation in the spermatheca (13:33 min), KCA-1 disperses from the vesicles. Yolk granules pack inward (12:16 min) while KCA-1 is on these vesicles. Time 0 is NEBD. C’. Magnified inset from -8 min shows KCA-1 on vesicles. C”. Magnified inset from 19 min shows KCA-1 dispersed from vesicles. E and F. Before ovulation (-16:50 min), KLC-1::AID::GFP (grayscale) is on vesicles that co-localize with VIT-2::Halo (cyan). During ovulation in the spermatheca (20:44 min), KLC-1 disperses from the vesicles. Yolk granules pack inward (19:33 min) while KLC-1 is on these vesicles. Time 0 is NEBD. E’. Magnified inset from -16:50 min shows KCA-1 on vesicles. C”. Magnified inset from 25:16 min shows KLC-1 dispersed from vesicles. G. Distance from the edge of the metaphase I spindle to the nearest cortex. H. Single plane from a deconvolved z-stack showing co-localization of KCA-1::AID::GFP with VIT-2::Halo labeled yolk granules. I. Fluorescence intensity of KCA-1 and yolk granules along the yellow line in H. J. *rab-7(RNAi)* makes yolk granules larger and makes KCA-1 appear as a ring on the outside of the yolk granules. K. Fluorescence intensity KCA-1 and yolk granules along the yellow line in I. Bar = 10 μm. **** p<.0001 Student’s t test.

In some control oocytes, the nucleus was positioned close to the cortex furthest from the spermatheca at NEBD so the spindle migrated only a short distance (Fig. 2A, 2D). However, in other cases the nucleus was centered at NEBD so that the spindle migrated a greater distance (Fig. 2F). In these cases, yolk granules could clearly be observed clearing out from the space between the spindle and cortex. The localization of KCA-1 and KLC-1 on yolk granules during spindle migration and the simultaneous timing of inward yolk granule movement and outward spindle movement are both consistent with the hypothesis that organelle packing causes spindle migration.

### Tailless Kinesin bound to Mitochondria is sufficient to restore spindle localization to the cortex during metaphase I

Because the cargo-binding C-terminal tail of kinesin-1 has been reported to bind many different cargos ^15^, it is possible that outward spindle movement is mediated by a subpopulation of kinesin that is not on yolk granules and might be obscured by the bright labeling of kinesin on yolk granules. To test whether inward packing of a single organelle is sufficient to move the spindle outward, we generated a mitochondria-only kinesin and tested its ability to restore outward spindle movement to *unc-116(g-null)* worms. To circumvent potential dominant lethality of attaching kinesin directly to mitochondria, we expressed two chimaeras in the *unc-116(g-null)* background (Fig. 3A). The first 420 amino acids of kinesin-1 heavy chain (K420) have motor activity but lack autoinhibition and cargo-binding domains ^23^. SSPB is a protein that has an increased affinity for iLID when exposed to 488 nm light ^24^. mKate is a red fluorescent protein ^25^.

**Figure 3.**
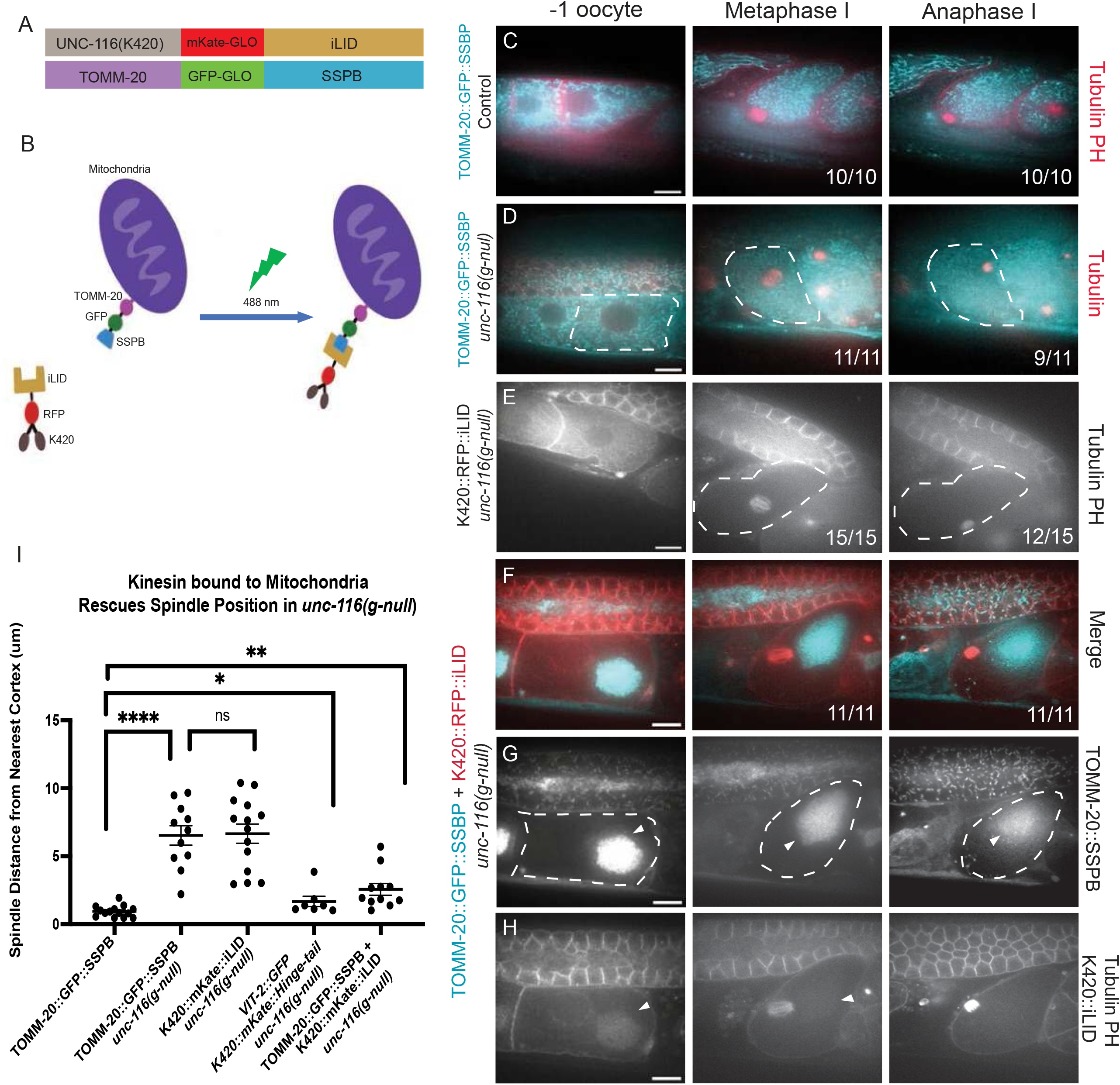
Tailless kinesin bound to mitochondria is sufficient to restore cortical spindle positioning in *unc-116(g-null)* worms. A. Transgene constructs used for optogenetically attaching kinesin motor domains to mitochondria. K420 (amino acids 1-420 of UNC-116 encoding just motor domains), mKate-GLO (red fluorescent protein) and iLID (improved light induced dimer). TOMM-20 (mitochondrial outer membrane protein, GFP-GLO (green fluorescent protein), and SSPB (binds iLID in blue light). B. Cartoon illustrating light-induced binding of kinesin motor domains to mitochondria. C. TOMM-20::GFP::SSPB labelled mitochondria (cyan) are evenly distributed in -1 oocytes, then packed inward at metaphase I in 10/10 control meiotic embryos. D. TOMM-20::GFP::SSPB labelled mitochondria remain evenly dispersed at metaphase I in *unc-116(g-null)* embryos. White dashed line highlights the cortex. The metaphase spindle is centered (11/11 embryos). During anaphase I, 9/11 meiotic spindles reached the cortex, while 2/11 remained in the middle. E. 15/15 metaphase I spindles were centered in *unc-116(g-null)* embryos expressing K420::mKate::iLID alone. During anaphase I, 12/15 spindles went to the cortex late. F-H. Combination of TOMM- 20::GFP::SSPB and K420::RFP::iLID leads to prematurely packed mitochondria in the oocyte and rescue of metaphase I spindle position in 11/11 *unc-116(g-null)* embryos. I. Distance from the edge of the metaphase I spindle to the nearest cortex. Bar = 10 μm. ns = not significant, ** p<.01, *** p<.001, **** p<.0001 Students t test.

TOMM-20 is a mitochondrial outer membrane protein with topology suitable for optogenetic coupling ^26^. Thus K420::mKate::iLID is expected to be recruited to the surface of mitochondria with TOMM-20::GFP::SSPB (Fig. 3B). Neither TOMM- 20::GFP::SSPB alone (Fig. 3C, 3D, Video S3) nor K420::mKate::iLID alone (Fig. 3C, 3E, Video S4) rescued the metaphase I spindle positioning defect of *unc-116(g-null)*. However, co-expression of both chimaeras resulted in co-localization of K420 with mitochondria in a large ball in the middle of oocytes and rescue of cortical spindle positioning (Fig. 3F-3I, Video S5). Rescue of spindle position by the mitochondria-only kinesin was comparable to rescue by a transgenic chimaera with the wild-type UNC-116 tail, K420::mKate::hinge-tail (Fig. 3I). This result suggests that inward packing of mitochondria is sufficient to move the spindle outward.

*unc-116(g-null)* strains expressing only TOMM-20::SSPB or only K420::iLID had higher frequencies of late translocation (Fig. 3D, 3E) and higher hatch rates (Table S1) than other *unc-116(g-null)* strains suggesting the formal possibility that the *duIs1* somatic rescuing transgene might be partially desilenced in these strains. However, depletion of cytoplasmic dynein heavy chain [*dhc-1(RNAi)*] prevented late translocation in these strains (Fig. S2), confirming that the late translocation is due to APC-activated dynein rather than residual UNC-116 activity. In addition, yolk granules were not packed inward, and instead were pushed outward in TOMM-20::GFP::SSPB/K420::mKate::iLID rescued *unc-116(g-null)* worms (Fig. S3A-F). These results support the interpretation that inward packing of only mitochondria can rescue outward spindle translocation in the complete absence of endogenous kinesin heavy chain.

A majority of *C. elegans* oocyte nuclei have been reported to migrate to the cortex furthest from the future site of fertilization by NEBD ^9^ so that the spindle most often migrates only a short distance ^5^. This nuclear migration has been reported to be kinesin-dependent in *C. elegans* ^13^. Thus there is the formal possibility that failed spindle migration is an indirect consequence of failed nuclear migration and the corresponding possibility that K420::iLID/TOMM-20::SSPB rescues nuclear migration rather than spindle migration. However, nuclear migration does not always occur in wild-type *C. elegans* ^9^(Fig. 2F; Fig S3J), but the spindle always migrates to the cortex. There are also cases of cortical nuclear position in *unc-116(g-null)* worms where the spindle then moves away from the cortex to the center of the embryo (Fig. S3J). These results support the idea that UNC-116 is required for spindle migration independent of nuclear migration and that K420::iLID/TOMM-20::SSPB rescues spindle position independently of nuclear migration.

### Tailless kinesin bound to LMP-1, TMEM-147, or EBP-2 is not sufficient to restore spindle localization to the cortex during metaphase I

To test whether attaching K420 minimal kinesin motor domains to other organelles is sufficient to move the spindle outward, we fused GFP::SSPB to LMP1 (lysosomal membrane protein), the endoplasmic reticulum (ER) protein, TMEM-147, or the microtubule plus end binding protein, EBP2. When co-expressed with K420::mKate::iLID, none of these chimaeras rescued the spindle positioning defect of *unc-116(g-null)* (Fig. 4A-4N). However, none of these fusions packed all of the targeted organelle into a large central ball as well as the TOMM-20 fusion packed mitochondria (Fig. 3). The TOMM-20/K420 pair packed essentially all of the mitochondria in oocytes (Fig. S4A, F) and after ovulation in meiotic embryos (Fig. 3). Packing of lysosomes by the LMP-1/K420 pair was much more variable. In some cases the ball started to form in the oocyte but dispersed during ovulation (Fig. 4D-E) and oocytes with the most pronounced ball of lysosomes (Fig. S4G, H) failed to ovulate. The TMEM-147/K420 pair sometimes formed a large ball of ER that persisted through ovulation (Fig. 4H-I), however, metaphase spindles are normally surrounded by an envelope of ER ^27^ and this envelope persisted in TMEM-147/K420 embryos (Fig. 4H-I). Thus K420 might move the spindle away from the cortex as it packs a different subset of ER than the ER that surrounds the meiotic spindle into a ball. The EBP-2/K420 pair was designed to drive microtubule-microtubule sliding which has been suggested to be how kinesin-1 carries out oocyte functions in *Drosophila* ^28^. The EBP-2/K420 pair generated a small ball of presumably cross-linked microtubules in metaphase embryos but did not rescue spindle positioning (Fig. 4K-N).

**Figure 4.**
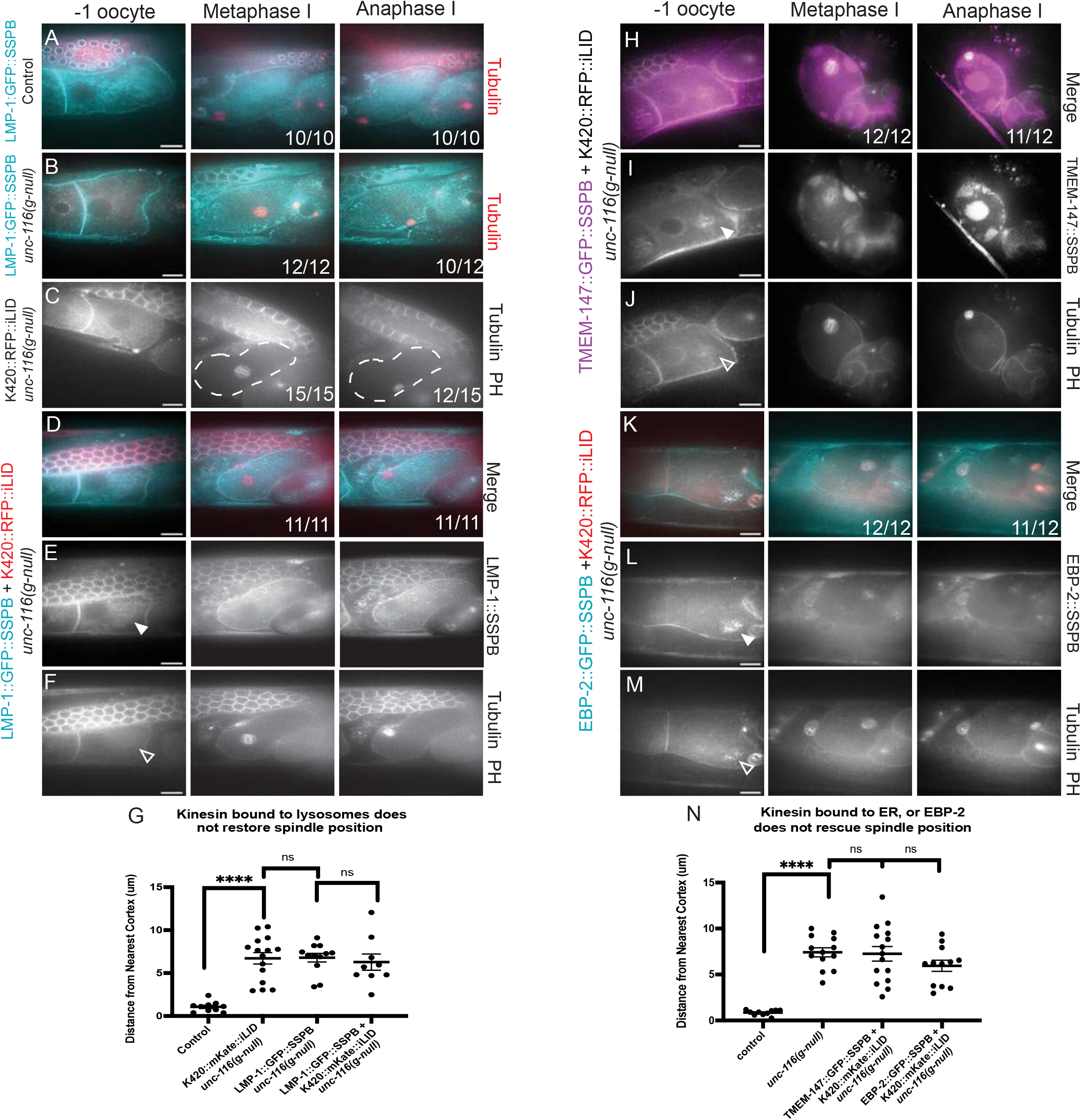
Tailless Kinesin bound to LMP-1, ER, or EBP-2 is not sufficient to restore spindle localization. A. LMP-1::GFP::SSPB (cyan) labels the plasma membrane and evenly dispersed vesicles in -1 oocytes of 10/10 control worms and these vesicles pack inward by metaphase I. B. LMP-1::GFP::SSPB vesicles did not pack inward in 12/12 *unc-116(g-null)* embryos and 12/12 had centered metaphase I spindles. C. K420::RFP::iLID alone did not rescue the centered metaphase I spindle in 15/15 *unc-116(g-null)* embryos. A white dashed line outlines the plasma membrane of the meiotic embryo. D-F. Combined expression of LMP-1::GFP::SSPB and K420::RFP::iLID did not rescue the centered metaphase I spindles in 11/11 *unc-116(g- null)* embryos. G. Distance from the edge of the metaphase I spindle to the nearest cortex. H-J. Combined expression of TMEM-147(ER)::GFP::SSPB and K420::mKate::iLID did not rescue the centered metaphase I spindles of 12/12 of *unc- 116(g-null embryos)*. An inward packed ball of ER is obvious in the anaphase I panel but is in a different focal plane at metaphase I. K-M. Combined expression of EBP- 2::GFP::SSPB and K420::RFP::iLID causes packing of presumably cross-linked microtubules in the -1 oocyte but did not rescue the centered metaphase I spindles of 12/12 *unc-116(g-null)* embryos. N. Distance from the edge of the metaphase I spindle to the nearest cortex. Bar = 10 μm. ns not significant, **** p< .0001 Students t test.

### Constitutively active kinesin packs yolk granules prematurely into a ball in -1 oocytes and pushes ER out of this ball

The localization of KCA-1 and KLC-1 on the surface of yolk granules suggested that packing of yolk granules might be the primary driver of kinesin-dependent outward spindle movement. Because there are no known membrane proteins specific to yolk granules, we were not able to target K420 specifically to yolk granules. To further investigate the normal cargo of kinesin during meiosis, we constructed a constitutively active kinesin with a normal cargo binding tail. K420::mKate::hinge tail has the C-terminal 246 aa of UNC-116 (aa 570-815) that contain light chain binding, cargo binding, and autoinhibitory domains. We hypothesized that the insertion of mKate would sterically block autoinhibitory interactions between the head and tail ^23^ thus generating a constitutively active kinesin. Yolk granules and ER were evenly distributed in control -1 oocytes (Fig. 5A). Expression of K420::mKate::hinge-tail in the presence of endogenous kinesin generated a large ball of yolk granules that excluded ER in -1 oocytes (Fig. 5B). This packing of yolk granules and exclusion of ER was more pronounced in the absence of endogenous kinesin (Fig. 5C). Premature packing of yolk granules by K420::mKate::hinge-tail supports the idea that this motor is constitutively active. The exclusion of ER from the packed ball of yolk granules supports the hypothesis that in wild-type, inward packing of yolk granules pushes on the sheet-like ER that envelopes the metaphase I spindle ^27^ by volume exclusion. Depletion of KCA-1 by RNAi prevented inward packing of yolk granules even though the K420::mKate::hinge-tail still packed into a ball (Fig. 5D). This result suggests that KCA- 1 is required for kinesin binding to yolk granules but is not required for kinesin’s motor activity. Constitutively active kinesin, in the presence (Fig. 5E-F) or absence (Fig. 5G) of KCA-1 did not pack mitochondria into the ball. This result suggests that yolk granules are not tethered to mitochondria during prophase. Overall our results show that inward packing of mitochondria is sufficient to move the ER-encased metaphase spindle outward and suggest that in wild-type, inward packing of yolk granules is the primary driver of kinesin-dependent outward spindle movement (Fig. 5H).

**Figure 5.**
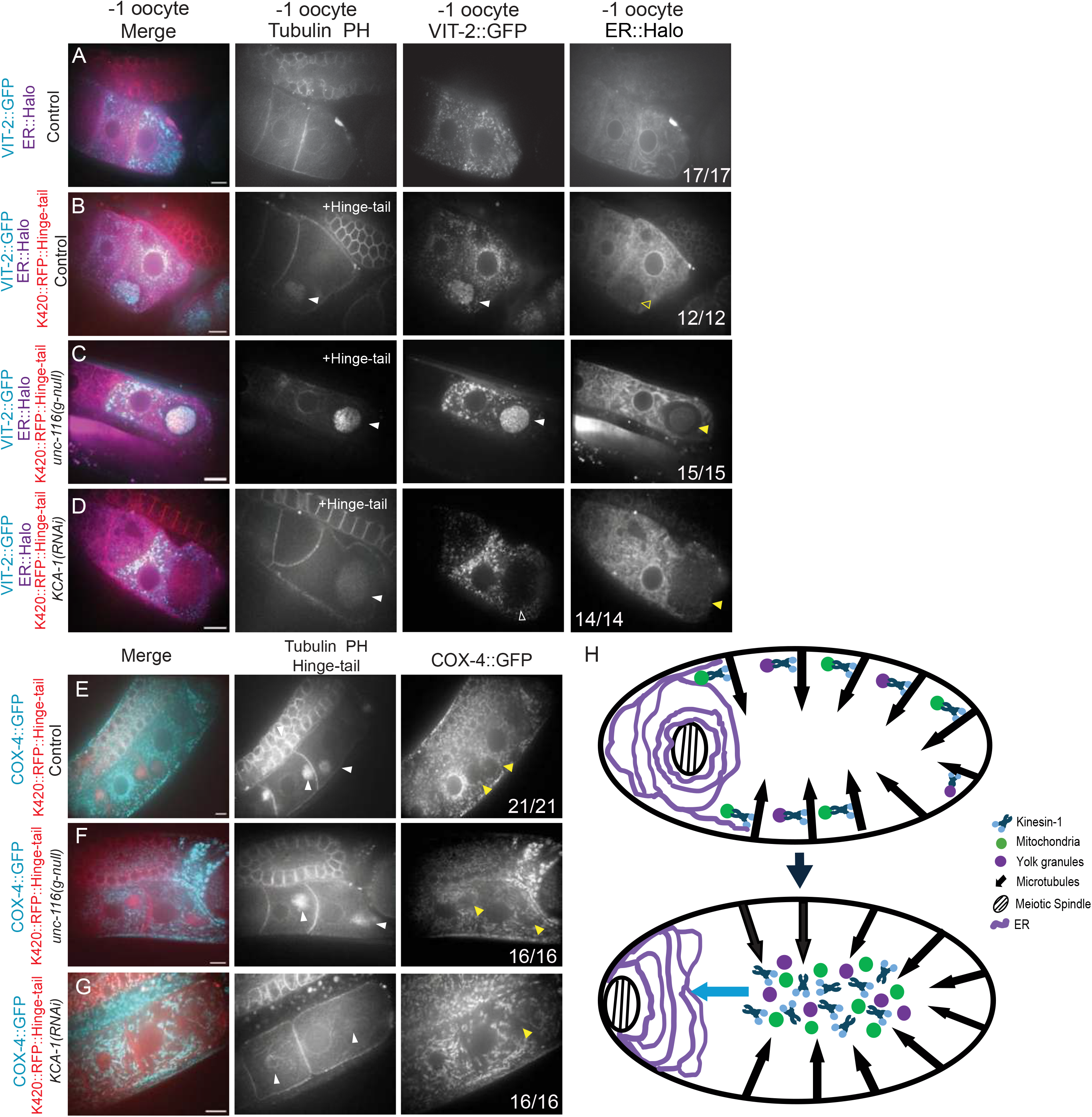
Constitutively active kinesin packs yolk granules inward prematurely in **-1 oocytes into a ball that pushes ER outward and KCA-1 is required to attach yolk granules to kinesin.** A. VIT-2-labeled yolk granules and ER are evenly dispersed in -1 oocytes of control worms. B-C. Expression of K420::mKate::Hinge-tail in control (B) or *unc-116(g-null)* worms causes yolk granules to pack into a ball that excludes ER. D. *kca-1(RNAi)* prevents yolk granule packing by K420::mKate::hinge tail. However K420::mKate::hinge tail still packs into a ball in the absence of KCA-1 and still excludes ER. E-G. The inwardly packed ball of K420::mKate::hinge tail excludes COX-4- labelled mitochondria in control (e), *unc-116(g-null)* oocytes (F) or *kca-1(RNAi)* (G). H. Model showing how Kinesin-1/ kinesin light chain/KCA-1 complex transports mitochondria and yolk granules towards the center of the meiotic embryo. This generates a force on the sheet-like ER that pushes the meiotic spindle to the cortex. Bar = 10 μm.

## Discussion

Our results support a model in which kinesin heavy chain/light chain/KCA-1 complexes transport yolk granules and mitochondria inward during NEBD and ovulation. We hypothesize that this transport is directed toward the plus ends of microtubules with minus ends at the cortex. In this model, the resulting packing of organelles generates an outward pushing force on the sheet-like ER that envelopes the prometaphase I meiotic spindle ^27^ by volume exclusion (Fig. 5H). Both mitochondria ^14^(Fig. 1) and yolk granules ^13^ are normally packed inward in a kinesin/KCA-1 dependent manner but inward packing of only mitochondria is sufficient to move the spindle outward (Fig. 3). Premature activation of kinesin before NEBD results in premature packing of yolk granules into a large ball that excludes ER (Fig. 5), supporting the idea that inward packing of organelles can push on the ER envelope surrounding the spindle. The timing of inward packing of mitochondria and yolk granules during NEBD, at the same time that the spindle is moved outward, also supports this model. KCA-1 and KLC-1 are localized on the surface of yolk granules before and during spindle translocation, also supporting the model. This localization was missed in an earlier study of metaphase I embryos ^13^ because KLC-1/KCA-1 dissociate from yolk granules after spindle translocation (Fig. 2). Yolk granules and mitochondria remain packed inward long after KLC-1/KCA-1 dissociate from yolk granules, suggesting that organelle positioning is maintained after active transport ceases, possibly due to the high viscosity of the cytoplasm.

Mouse meiotic spindle translocation depends on actin and formin 2 ^3^ rather than microtubules and kinesin, however, similar physical mechanisms could be at work.

Mouse oocytes do not have yolk granules but have abundant mitochondria that are packed inward at the time the metaphase I spindle moves outward. Duan et al. proposed a model in which formin 2 on ER-associated vesicles around the spindle nucleates actin filaments that push on the mitochondria ball in the middle of the oocyte. Dominant expression of the myosinVB tail or depletion of formin 2 both dispersed the mitochondria and blocked spindle translocation, supporting the model ^29^. However, a different study found that depletion of ZAR1 dispersed the mitochondrial ball without apparently affecting spindle translocation ^30^. Thus the role of inward mitochondrial packing in mouse meiotic spindle translocation remains unclear. Yolk granules are also packed inward in the oocytes of zebrafish ^31, 32^ and leech ^33^. All of these species undergo fertilization-dependent cortical granule exocytosis. Thus clearing of yolk granules and/or mitochondria from the cortex might be important in preventing interference with cortical granule exocytosis, independently of spindle translocation.

Defective cortical granule exocytosis observed in *kca-1(RNAi)* embryos ^34^ supports this idea in *C. elegans*. Although mammals do not have vitellogenin-containing yolk granules, KCA-1 is present on vesicles before endocytosis of vitellogenin (Fig. S1G). Identification of other molecules on the surface of these vesicles will be important in defining how KCA-1 is targeted and will be important in understanding if this class of vesicles exists in mammalian oocytes.

Microtubules with minus ends attached to the cell cortex is a cytoskeletal organization reported in *Drosophila* oocytes ^35, 36^, *Xenopus* oocytes ^37^, and in polarized epithelial cells of *C. elegans* ^38^ and mammals ^39^. In *Drosophila* oocytes, a polarized gradient of cortical microtubule density has been suggested to mediate preferential kinesin-dependent transport of oskar RNA away from the cortex at one end of the oocyte so that it accumulates at the end with lowest microtubule density ^40^. In polarized epithelial cells, the density of microtubule minus ends at the cortex is highest at the apical plasma membrane such that plus ends extend toward the basolateral surface ^38,39^. Cortical localization of microtubule minus-end associated proteins like γ-tubulin, patronin/CAMSAP, or ninein is often used to infer this microtubule arrangement ^41^. ψ-tubulin is concentrated at the plasma membrane of earlier *C. elegans* oocytes ^42^ but has not been detected at the plasma membrane of -1 oocytes or meiotic embryos ^43, 44^ suggesting the possibility of an unidentified minus end anchoring protein at these stages. Microtubule minus ends at the cortex have not been reported in mouse oocytes. Instead, a cytoplasmic network of microtubules has been reported to restrain actin-driven spindle movement ^45^ and the plus ends of astral microtubules have been reported to interact with the cortex after the spindle has moved close to the cortex ^46^. This latter interaction may be analogous to the APC and dynein-dependent spindle rotation reported in *C. elegans* ^5–7, 10^.

## Materials and Methods

### *C. elegans* strains

Complete genotypes of C. elegans strains used in this study are in Table S2.

Hatch rates for each strain are in Table S1. Sequences of transgenes are in Data S1.

### CRISPR editing

AID::GFP tags were added to the endogenous *kca-1* and *klc-1* loci and HALOtag was added to the *vit-2* locus by SUNY Biotech. Sequences of insertions are in Data S1.

### Transgenes

Optogenetic chimaeras were assembled with germline-optimized ^47^ versions of GFP, mKate, SSPB and iLID, and PCR amplified genomic regions of UNC-116, TOMM- 20, LAMP-1, TMEM-147(ZK418.5), EBP-2, *mex-5* promoter and *tbb-2* 3’UTR in the RMCE vector pLF3FShC ^48^. Plasmids were injected by In vivo Biosystems using Inject Express service then screened as described ^48^. K420::mKate::iLID was inserted at the jsTi1490 chromosome IV landing site in NM5176. All SSPB transgenes were inserted at the jsSi1579 chromosome II landing site in NM5402 and FLP recombinase was removed by outcrossing.

### RNAi, auxin treatment, and HALO-646 labeling

*dhc-1(RNAi)* and *rab-7(RNAi)* were conducted by placing L4 worms on MYOB plates seeded with HT115 bacteria expressing dsRNA as described by ^49^ for 24 hrs. *kca-1(RNAi)* was conducted for 30 hrs. For auxin-mediated degradation, auxin (indole acetic acid) was added to molten MYOB agar from a 400mM stock solution in ethanol to a final concentration of 4mM auxin before pouring plates, which were subsequently seeded with OP50 bacteria. Young adult worms were placed on auxin plates for 3 hrs before anesthetizing and imaging. Janelia fluor 646 HaloTag Ligand was diluted from a 200 uM stock in DMSO into M9 buffer to a final concentration of 2.5 uM. 100 ul of this dilution was pipetted onto a lawn of OP50 bacteria on a 60 mm MYOB plate with 40-80 worms. Worms were labeled for 20-24 hrs before imaging.

### Microscopy and in utero time-lapse imaging

L4 larvae were incubated at 20°C overnight on MYOB plates seeded with OP50. Worms were anesthetized by picking young adult hermaphrodites (for the best imaging quality) into a solution of 0.1% tricaine, 0.01% tetramisole in PBS in a watch glass for 30min as previously described ^9, 50^. Worms were then transferred in a small volume to a thin agarose pad (2% agarose in water) on a slide. Additional anesthetic in PBS solution was pipetted around the edges of the agarose pad, and a 22-x-30-mm coverslip was place on top. Because *unc-116(g-null)* worms have a *rol* phenotype, which causes worms to twist around their long axis, these worms were mounted in grooves imprinted in the agarose pad with a Lana Del Rey vinyl record as described ^51^. Mounting in grooves partially unwound the twisted worms to improve imaging. The slide was inverted and placed on the stage of an inverted microscope. -1 oocytes or meiotic embryos were identified by brightfield microscopy before initiating time-lapse fluorescence. For all live imaging, the stage and immersion oil temperature was 20°C-24°C. For all time-lapse data, single-focal plane images were acquired with a Solamere spinning disk confocal microscope equipped with an Olympus IX-70 stand, Yokogawa CSU10, Hamamatsu ORCA-Quest qCMOS (quantitative complementary metal oxide semiconductor) detector, Olympus 100x/1.35 objective, 100-mW Coherent Obis laser set at 30% power, and MicroManager software control. Pixel size was 46nm. Exposures were 100ms for the ORCA-Quest qCMOS detector. Time interval between image pairs was 5 seconds. Focus was adjusted manually during time-lapse imaging. For single time point imaging of -1 oocytes, z stacks were collected at 1 um z intervals using a PI E662 LVPZT piezo objective collar.

### Optogenetic coupling

The TOMM-20/iLID pair formed a ball of mitochondria in oocytes even before 488 nm illumination (Fig. S4A, F). iLID pairs with LMP-1, TMEM-147, or EBP-2 only formed a ball of cargo in oocytes after 600 – 900 s of illumination with pulses of 488 nm light (200 ms exposures over 5 sec) (Fig. S4).

### Image Analysis

Spindle position in all meiotic embryos were determined using the line tool in Fiji. Measurements began from the edge of the spindle to the nearest cortex of the meiotic embryo. Student’s t test was performed in GraphPad to calculate P values (Fig. 1K-N, Fig. 2G, Fig. 3I, Fig. 4G & 4N, Fig. S3I). Nuclear position was also measured using the line tool in Fiji. Measurements began from the edge of the nucleus to the nearest cortex. Student’s t test was performed in GraphPad to calculate P values (Fig. S3J).

Fluorescence intensity was measured in Fiji using the line tool and measured with the Plot Profile tool. Measurements were then overlayed in GraphPad (Fig. 2I & 2K, Fig. S1B, S1D & S1F). Denoising of images with either KCA-1 or KLC-1 and mitochondria labeled were processed on ZEN Blue using the Intellisis Denoising program (Fig. S1A & S1E). Deconvolution of images with KCA-1 or KLC-1 and yolk granules labeling was done by using Huygens Essential x11 (Fig. 2H & 2J, Fig. S1C).

## Acknowledgements

We thank Dan Dickenson for germline-optimized versions of GFP, mKate, SSPB, and iLID. We thank the Caenorhabditis Genetics Center, which is funded by NIH Office of Research Infrastructure Programs (P40 OD010440), for strains. We thank Dan Starr for critical reading of the manuscript.

---

**Figure S1.**
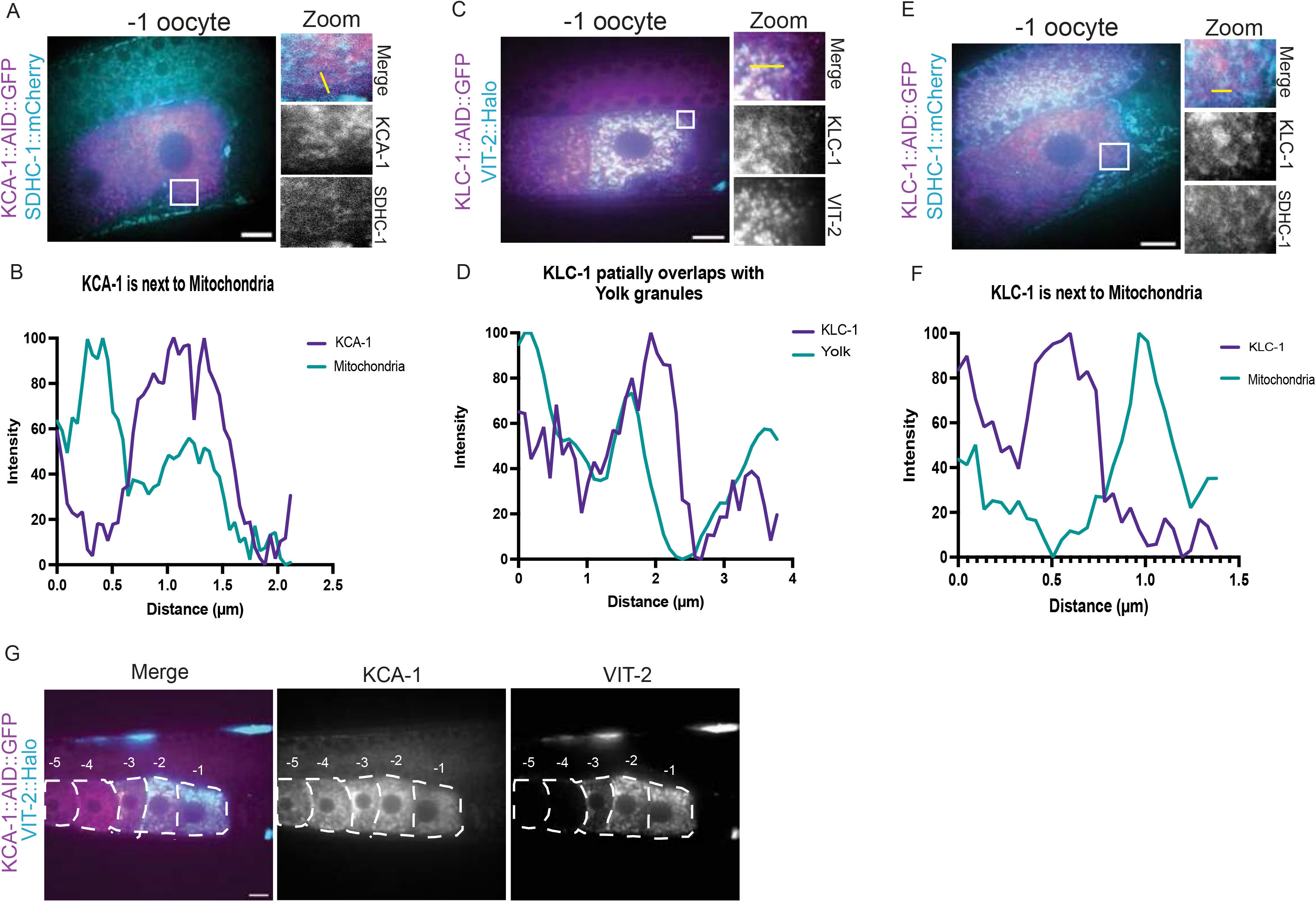
KCA-1 and KLC-1 are adjacent to mitochondria and KLC-1 co-localizes with yolk granules. Denoised single plane images from time-lapse sequences. A. KCA- 1::AID::GFP (magenta) does not co-localize with mitochondria (cyan). Zoom images are of the white square in the left image. B. Fluorescence intensity along the yellow line shown in A. C. KLC-1::AID::GFP (magenta) co-localizes with VIT-2-labeled yolk granules (cyan). Zoom images are of the white square in the left image. D. Fluorescence intensity along the yellow line in C. E. KLC-1::AID::GFP (magenta) does not co-localize with mitochondria (cyan). F. Fluorescence intensity along the yellow line in E. G. KCA-1::AID::GFP is present in the -1 through -5 oocytes, while yolk granules are only present in the -1 through -3 oocytes. Bar = 10 μm.

**Figure S2.**
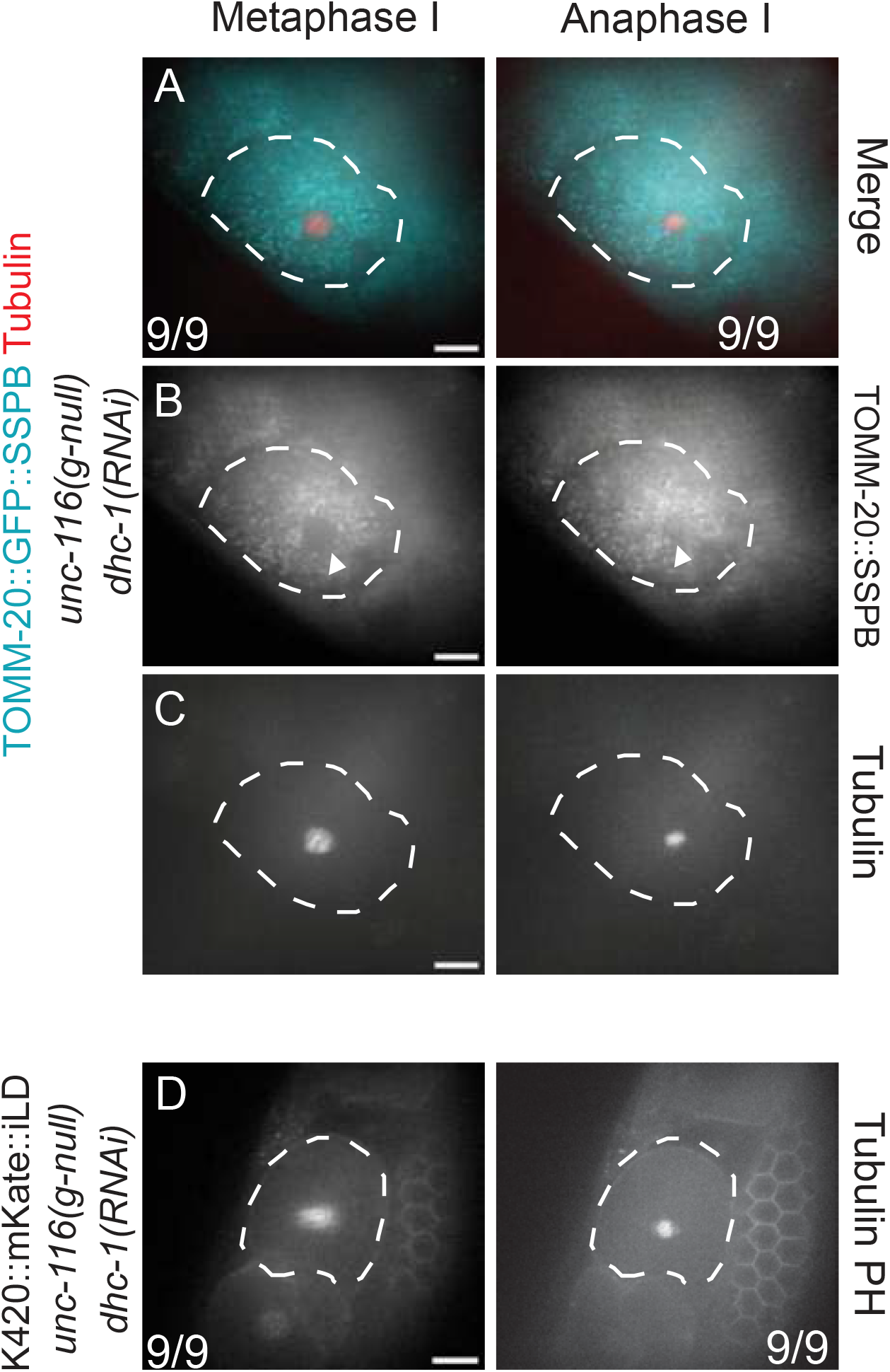
**Shortened anaphase spindles remain centered in the double mutation of *unc- 116(gk5722) duIs1 and dhc-1(RNAi).*** Single focal planes taken from time-lapse sequences. A. Merged images of the double mutation of *unc-116(gk5722) duIs1* and *dhc-1(RNAi)* during metaphase I and anaphase I in a strain expressing TOMM- 20::GFP::SSPB only. B. Grayscale of TOMM-20::GFP::SSPB. White arrowheads points to the empty space that is taken up by the spindle. C. Grayscale of the mKate::tubulin-labeled spindle. D. Grayscale images K420::mKate::iLID and mKate::tubulin in the double mutation of *unc-116(gk5722) duIs1* and *dhc-1(RNAi)* during metaphase I and anaphase I in a strain expressing K420::mKate::iLID only. White dashed boarder outlines the cortex of the meiotic embryo. Bar = 10 μm

**Figure S3.**
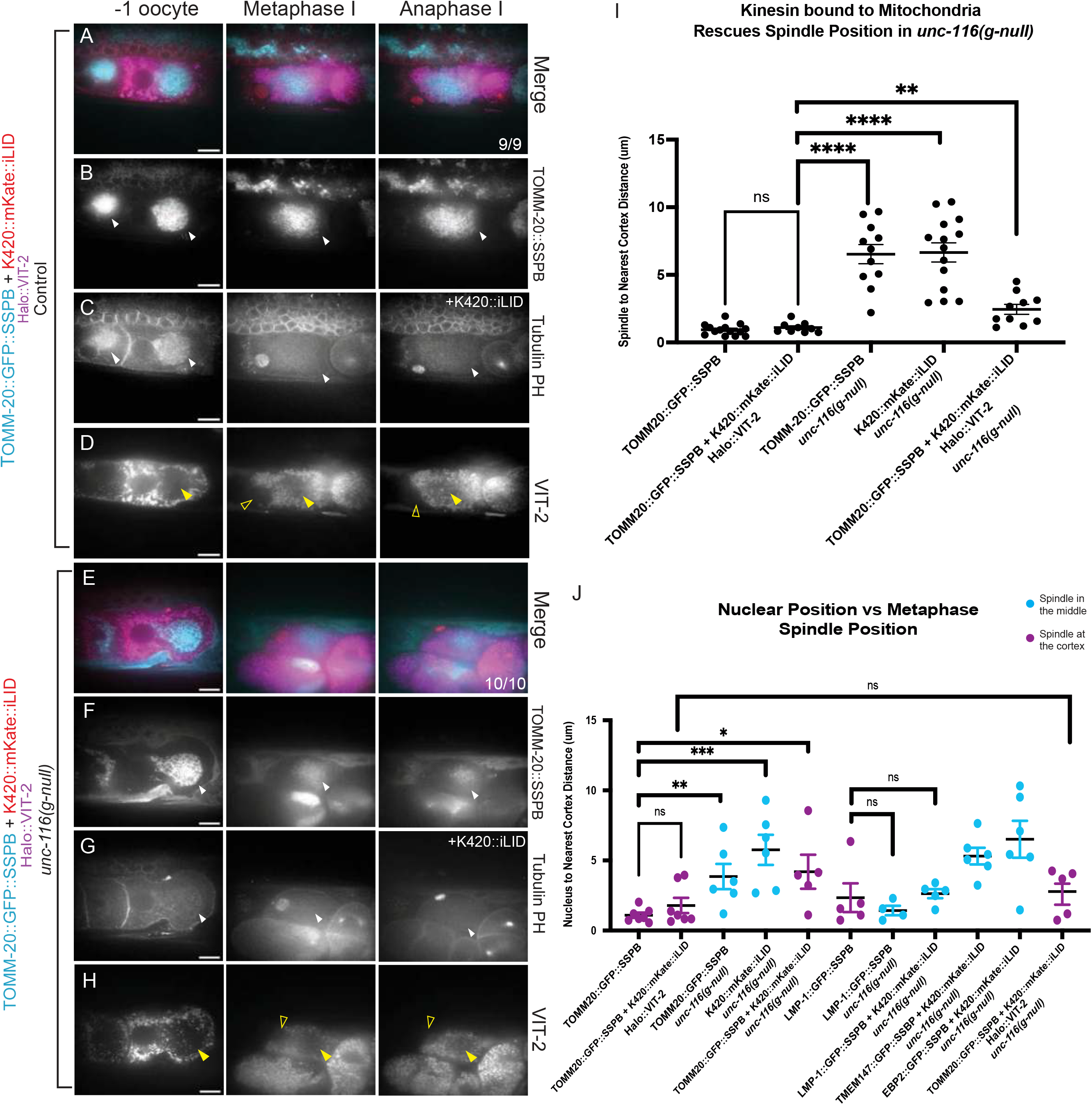
Tailless kinesin bound to mitochondria does not pack in yolk granules in the *unc-116(g-null)* background. Single focal plane images from time-lapse sequences. A – D. Optogenetic pairing of TOMM-20::GFP::SSPB with K420::mKate::iLID in a wild- type background results in a packed ball of mitochondria that excludes yolk granules. E - H. Optogenetic pairing of TOMM-20::GFP::SSPB with K420::mKate::iLID in the *unc- 116(g-null)* background results in a packed ball of mitochondria that excludes yolk granules. I. Distance from the edge of the metaphase I spindle to the nearest cortex. J. Distance from the edge of the nucleus to the nearest cortex at NEBD. Magenta symbols indicate oocytes in which the subsequent metaphase I spindle was at the cortex. Cyan symbols indicate oocytes in which the subsequent metaphase I spindle was centered. Ns = not significant; * p < .05; ** p < .01; *** p < .001; **** p < .0001 Students t test. Bar = 10 μm.

**Figure S4.**
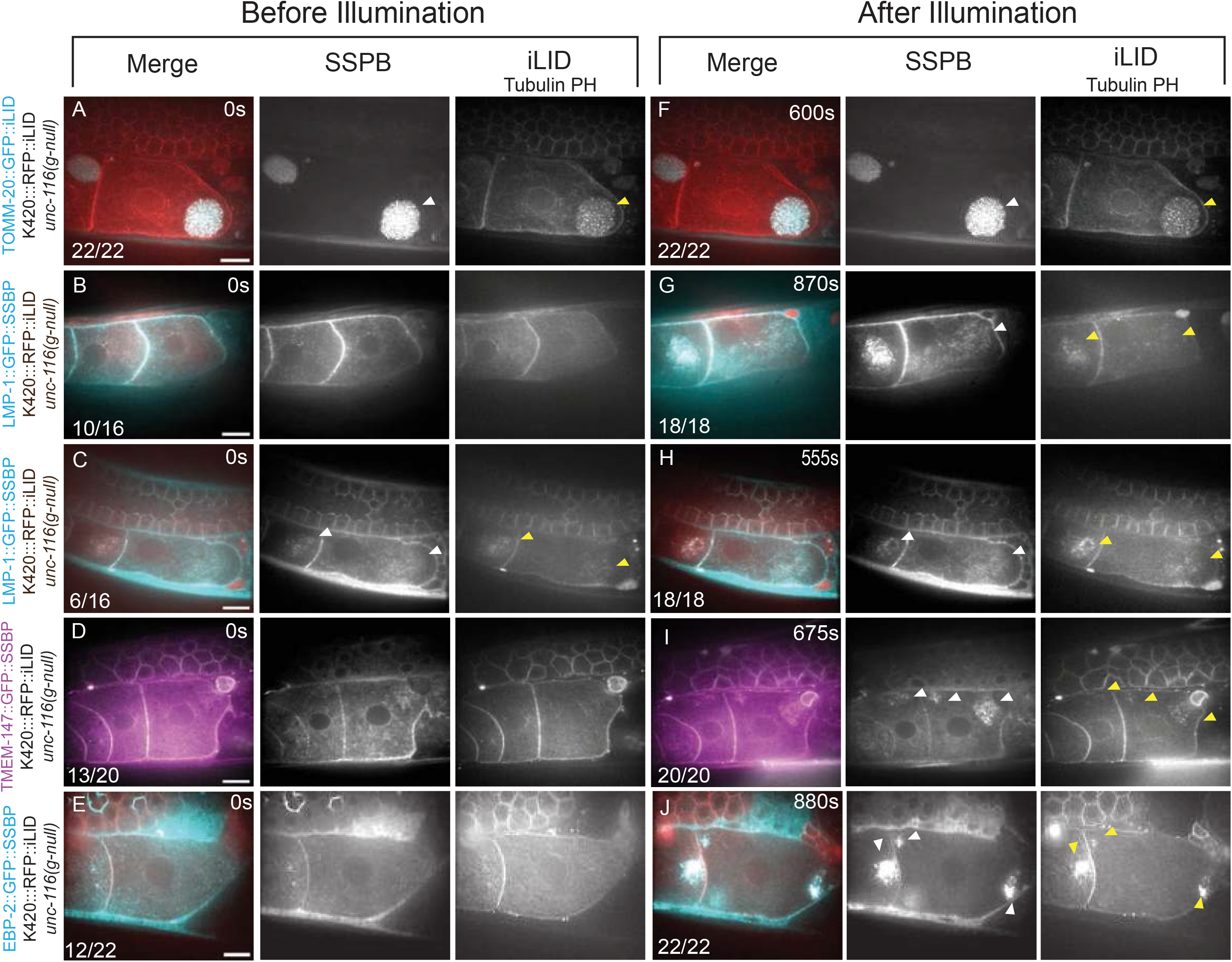
Before and after illumination of optogenetic pairings in oocytes. A-J. Single focal plane images from time-lapse sequences. “Before Illumination” indicates the first frame of a time-lapse sequence. The time indicated under “After Illumination” indicates the time required to generate a robust ball of cargo in that particular example. A. The TOMM-20/K420 pair packed all of the mitochondria into a ball before illumination in 22/22 worms. B and C. The LMP-1 /K420 pair did not pack lysosomes into a ball until after illumination in 10/16 worms but showed light-independent packing in 6/16 worms. D. The TMEM-147/K420 pair did not pack ER into a ball until after illumination in 13/20 worms but displayed light-independent packing in 7/20 worms. E. The EBP-2/ K420 pair did not pack microtubules into a ball until after illumination in 12/22 but light-independent packing was observed in 8/22 worms. F-J. All combinations showed packing of cargo into a ball after illumination. Many oocytes with the most dramatic balls did not ovulate. Bar = 10 μm.

## Video legends

**Video S1. Time-lapse video of KCA-1 and yolk granules during ovulation.** Meiosis I filmed in utero in a strain with endogenously tagged KCA-1::AID::GFP (greyscale) and endogenously tagged VIT-2::Halo (Yolk granules; cyan). mKate::tubulin (spindle) and mCherry:PH (plasma membrane) are labeled in red. Before ovulation KCA-1::AID::GFP is on vesicles that co-localize with yolk granules. During ovulation KCA-1::AID::GFP comes off the vesicles and appears to be on cytoplasmic filaments. Yolk granules stay packed after dissociation of KCA-1. Bar = 10 μm

**Video S2. Time-lapse video of KLC-1 and yolk granules during ovulation.** Meiosis I filmed in utero in a strain with endogenously tagged KLC-1::AID::GFP (greyscale) and endogenously tagged VIT-2::Halo (Yolk granules; cyan). mKate::tubulin (spindle) and mCherry:PH (plasma membrane) are labeled in red. Before ovulation KLC-1::AID::GFP is on vesicles that co-localize with yolk granules. During ovulation KLC-1::AID::GFP comes off the vesicles and appears to be on cytoplasmic filaments. Yolk granules stay packed after dissociation of KLC-1. Bar = 10 μm

**Video S3. Time-lapse video of TOMM-20::GFP::SSPB in the *unc-116(g-null)* background.** Meiosis I filmed in utero with TOMM-20::GFP::SSPB (mitochondria; cyan) in the *unc-116(g-null)* background. The meiotic spindle (red) is centered during metaphase I and reaches the cortex during anaphase I. Mitochondria are relaxed out toward the cortex. Bar = 10 μm

**Video S4. Time-lapse video of K420::mKate::iLID in *unc-116(g-null)* background.** Meiosis I filmed in utero with K420::mKate::iLID, mKate::tubulin (spindle), and mCherry::PH (plasma membrane) in the *unc-116(g-null)* background, all in grayscale. The meiotic spindle is centered during metaphase I and reaches the cortex during anaphase I. Bar = 10 μm

**Video S5. Time-lapse video of TOMM-20::GFP::SSPB and K420::mKate::iLID in the *unc-116(g-null)* background.** Meiosis I filmed in utero with the combination of TOMM- 20::GFP::SSPB (cyan) and K420::mKate::iLID (red) in the *unc-116(g-null)* background. Tubulin (spindle) and mCherry::PH (plasma membrane) are also in red. Mitochondria are packed into a tight ball at the future posterior end of the oocyte. During ovulation the mitochondria remain packed toward the center and the spindle is localized at the cortex during metaphase I and anaphase I. Bar = 10 μm

**Table S1:**
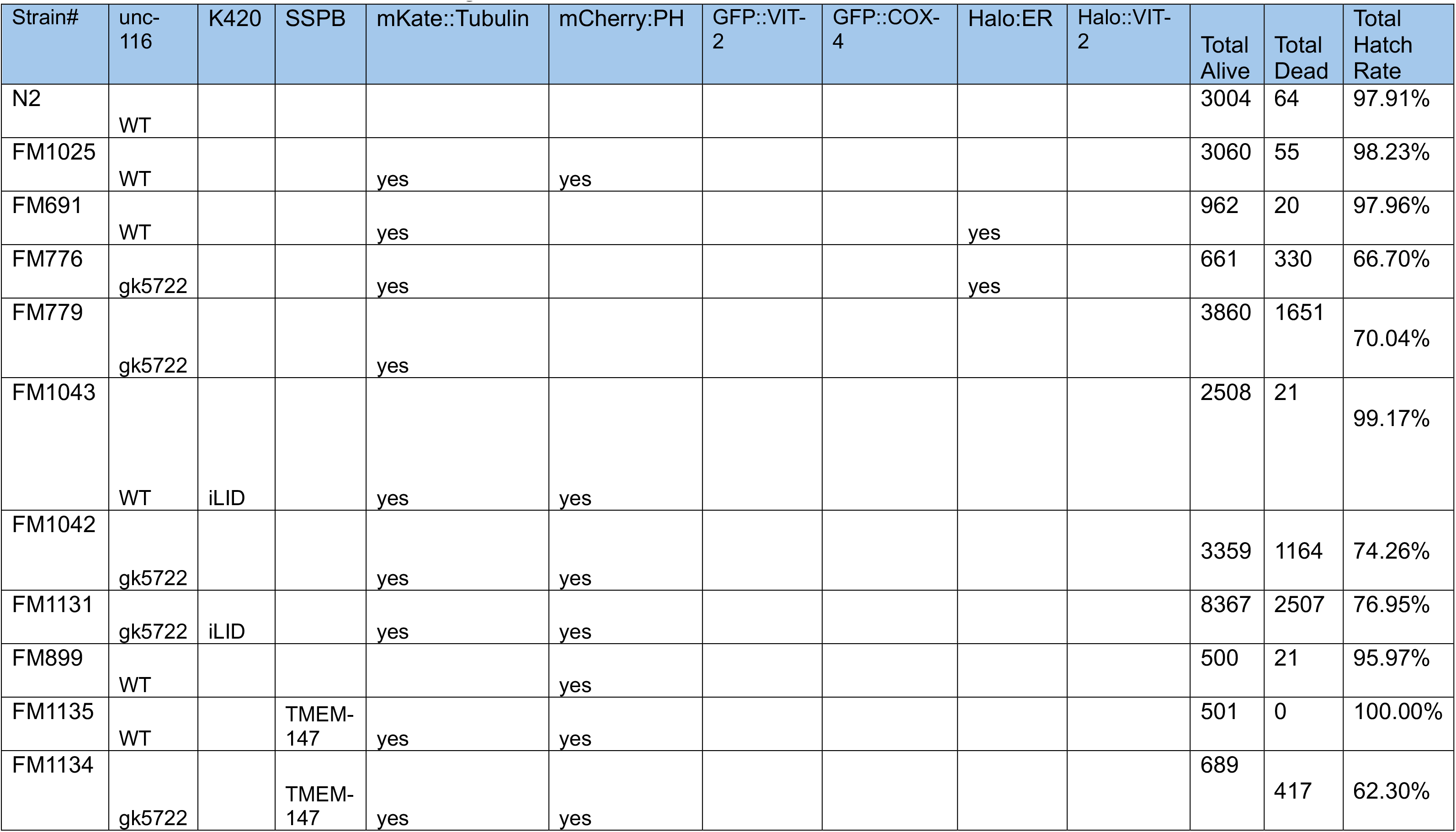
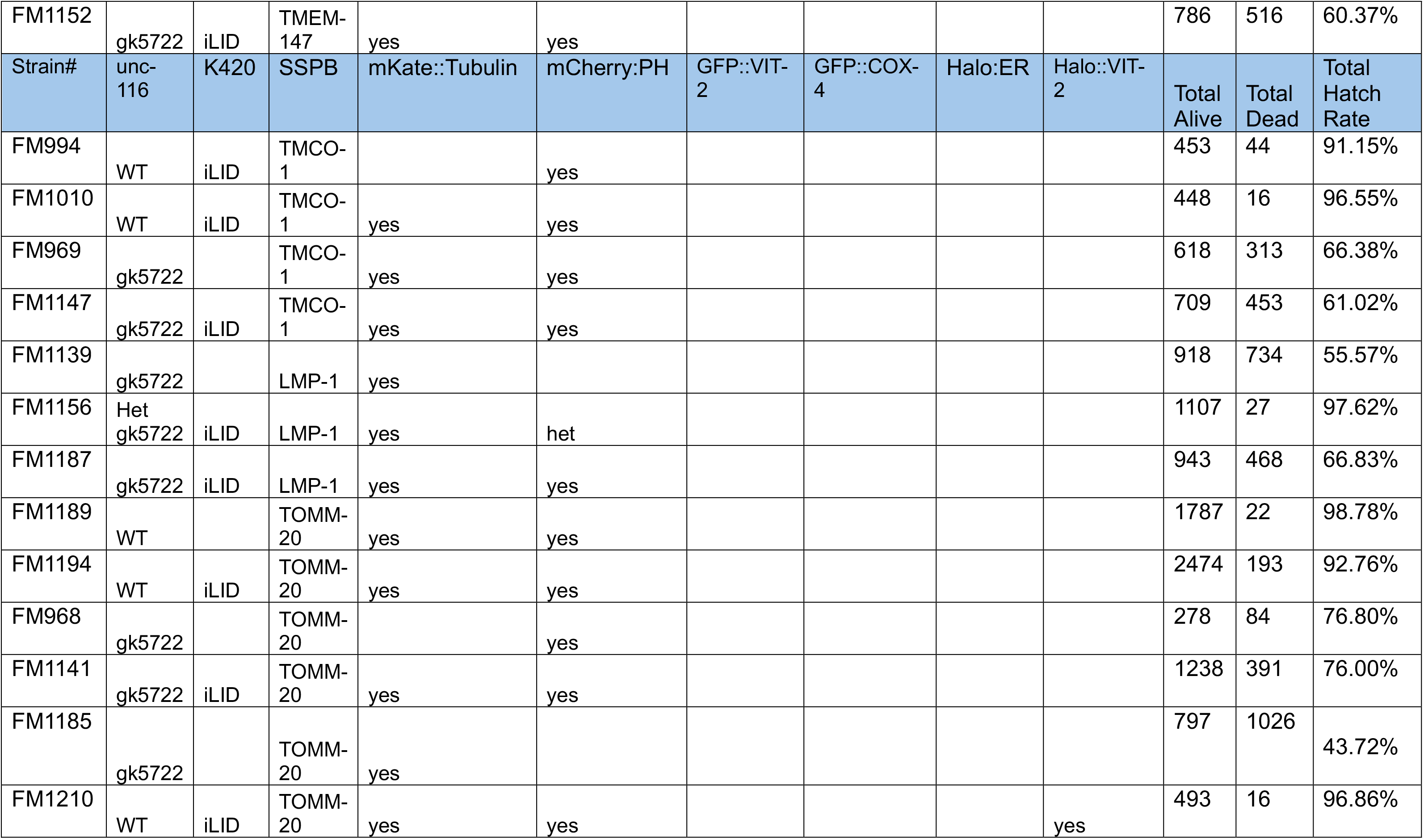
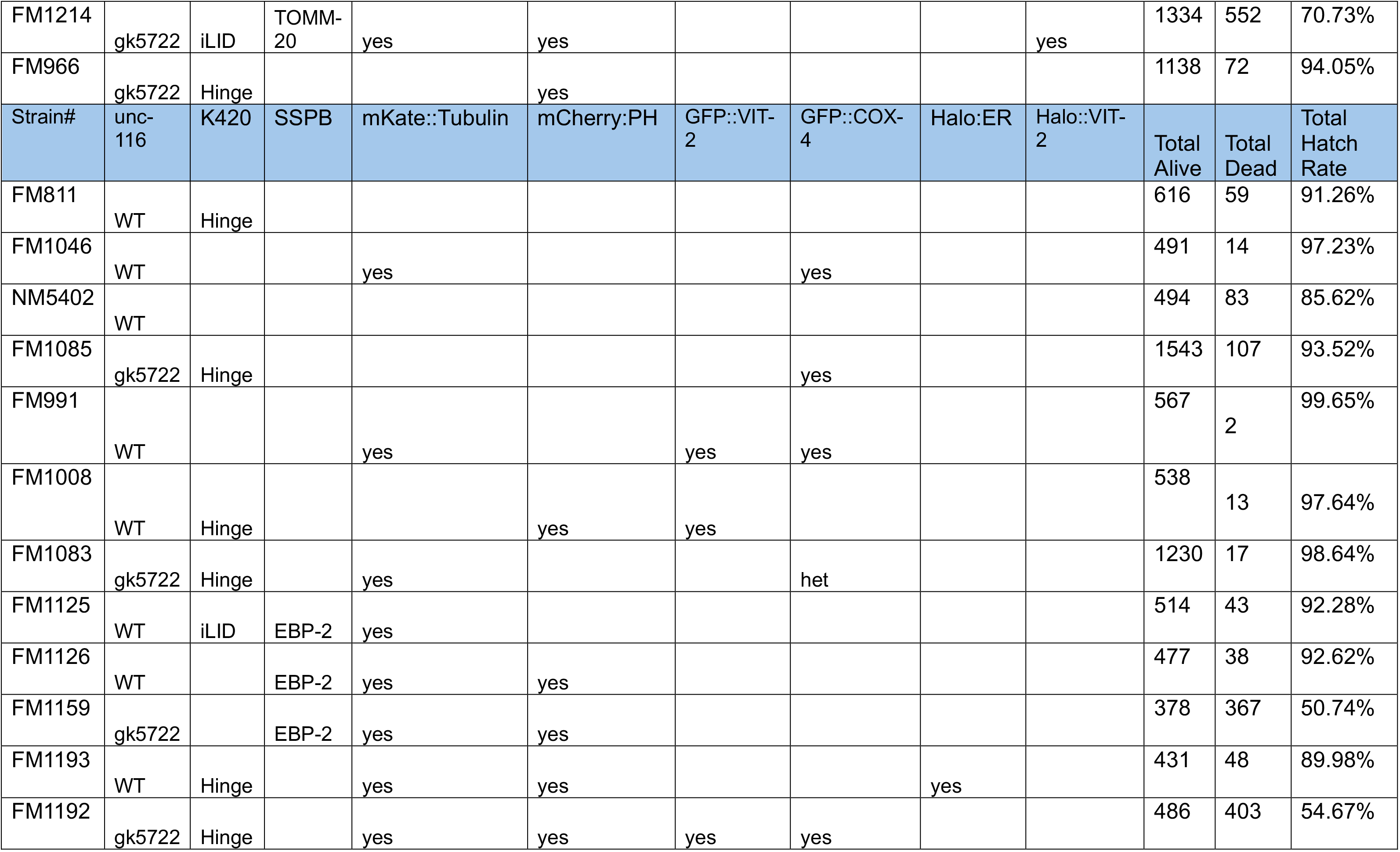
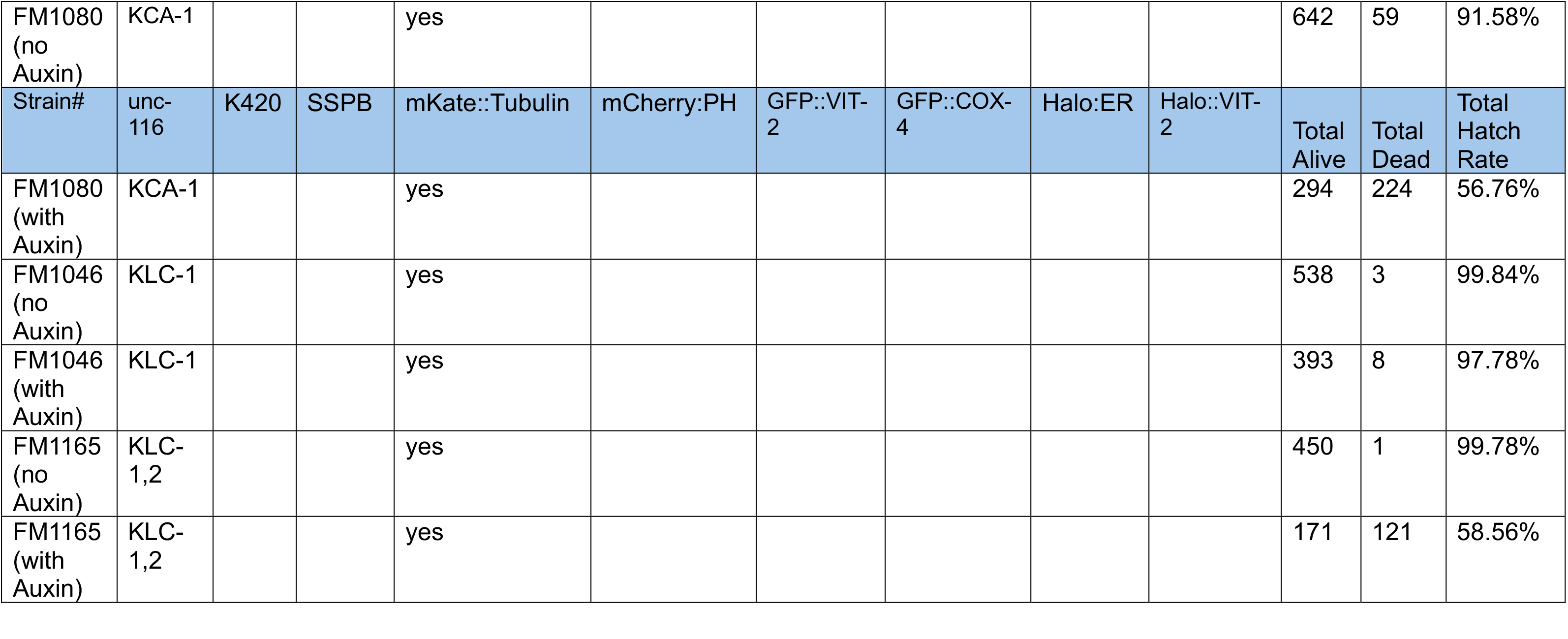
Hatch counts for all C. elegans strains.

| Strain# | unc-116 | K420 | SSPB | mKate::Tubulin | mCherry:PH | GFP::VIT-2 | GFP::COX-4 | Halo:ER | Halo::VIT-2 | Total Alive | Total Dead | Total Hatch Rate |
| --- | --- | --- | --- | --- | --- | --- | --- | --- | --- | --- | --- | --- |
| N2 | WT |  |  |  |  |  |  |  |  | 3004 | 64 | 97.91% |
| FM1025 | WT |  |  | yes | yes |  |  |  |  | 3060 | 55 | 98.23% |
| FM691 | WT |  |  | yes |  |  |  | yes |  | 962 | 20 | 97.96% |
| FM776 | gk5722 |  |  | yes |  |  |  | yes |  | 661 | 330 | 66.70% |
| FM779 | gk5722 |  |  | yes |  |  |  |  |  | 3860 | 1651 | 70.04% |
| FM1043 | WT | iLID |  | yes | yes |  |  |  |  | 2508 | 21 | 99.17% |
| FM1042 | gk5722 |  |  | yes | yes |  |  |  |  | 3359 | 1164 | 74.26% |
| FM1131 | gk5722 | iLID |  | yes | yes |  |  |  |  | 8367 | 2507 | 76.95% |
| FM899 | WT |  |  |  | yes |  |  |  |  | 500 | 21 | 95.97% |
| FM1135 | WT |  | TMEM-147 | yes | yes |  |  |  |  | 501 | 0 | 100.00% |
| FM1134 | gk5722 |  | TMEM-147 | yes | yes |  |  |  |  | 689 | 417 | 62.30% |
| FM1152 | gk5722 | iLID | TMEM-147 | yes | yes |  |  |  |  | 786 | 516 | 60.37% |
| Strain# | unc-116 | K420 | SSPB | mKate::Tubulin | mCherry:PH | GFP::VIT-2 | GFP::COX-4 | Halo:ER | Halo::VIT-2 | Total Alive | Total Dead | Total Hatch Rate |
| FM994 | WT | iLID | TMCO-1 |  | yes |  |  |  |  | 453 | 44 | 91.15% |
| FM1010 | WT | iLID | TMCO-1 | yes | yes |  |  |  |  | 448 | 16 | 96.55% |
| FM969 | gk5722 |  | TMCO-1 | yes | yes |  |  |  |  | 618 | 313 | 66.38% |
| FM1147 | gk5722 | iLID | TMCO-1 | yes | yes |  |  |  |  | 709 | 453 | 61.02% |
| FM1139 | gk5722 |  | LMP-1 | yes |  |  |  |  |  | 918 | 734 | 55.57% |
| FM1156 | Het gk5722 | iLID | LMP-1 | yes | het |  |  |  |  | 1107 | 27 | 97.62% |
| FM1187 | gk5722 | iLID | LMP-1 | yes | yes |  |  |  |  | 943 | 468 | 66.83% |
| FM1189 | WT |  | TOMM-20 | yes | yes |  |  |  |  | 1787 | 22 | 98.78% |
| FM1194 | WT | iLID | TOMM-20 | yes | yes |  |  |  |  | 2474 | 193 | 92.76% |
| FM968 | gk5722 |  | TOMM-20 |  | yes |  |  |  |  | 278 | 84 | 76.80% |
| FM1141 | gk5722 | iLID | TOMM-20 | yes | yes |  |  |  |  | 1238 | 391 | 76.00% |
| FM1185 | gk5722 |  | TOMM-20 | yes |  |  |  |  |  | 797 | 1026 | 43.72% |
| FM1210 | WT | iLID | TOMM-20 | yes | yes |  |  |  | yes | 493 | 16 | 96.86% |
| FM1214 | gk5722 | iLID | TOMM-20 | yes | yes |  |  |  | yes | 1334 | 552 | 70.73% |
| FM966 | gk5722 | Hinge |  |  | yes |  |  |  |  | 1138 | 72 | 94.05% |
| Strain# | unc-116 | K420 | SSPB | mKate::Tubulin | mCherry:PH | GFP::VIT-2 | GFP::COX-4 | Halo:ER | Halo::VIT-2 | Total Alive | Total Dead | Total Hatch Rate |
| FM811 | WT | Hinge |  |  |  |  |  |  |  | 616 | 59 | 91.26% |
| FM1046 | WT |  |  | yes |  |  | yes |  |  | 491 | 14 | 97.23% |
| NM5402 | WT |  |  |  |  |  |  |  |  | 494 | 83 | 85.62% |
| FM1085 | gk5722 | Hinge |  |  |  |  | yes |  |  | 1543 | 107 | 93.52% |
| FM991 | WT |  |  | yes |  | yes | yes |  |  | 567 | 2 | 99.65% |
| FM1008 | WT | Hinge |  |  | yes | yes |  |  |  | 538 | 13 | 97.64% |
| FM1083 | gk5722 | Hinge |  | yes |  |  | het |  |  | 1230 | 17 | 98.64% |
| FM1125 | WT | iLID | EBP-2 | yes |  |  |  |  |  | 514 | 43 | 92.28% |
| FM1126 | WT |  | EBP-2 | yes | yes |  |  |  |  | 477 | 38 | 92.62% |
| FM1159 | gk5722 |  | EBP-2 | yes | yes |  |  |  |  | 378 | 367 | 50.74% |
| FM1193 | WT | Hinge |  | yes | yes |  |  | yes |  | 431 | 48 | 89.98% |
| FM1192 | gk5722 | Hinge |  | yes | yes | yes | yes |  |  | 486 | 403 | 54.67% |
| FM1080<br>(no<br>Auxin) | KCA-1 |  |  | yes |  |  |  |  |  | 642 | 59 | 91.58% |
| Strain# | unc-116 | K420 | SSPB | mKate::Tubulin | mCherry:PH | GFP::VIT-2 | GFP::COX-4 | Halo:ER | Halo::VIT-2 | Total<br>Alive | Total<br>Dead | Total<br>Hatch<br>Rate |
| FM1080<br>(with<br>Auxin) | KCA-1 |  |  | yes |  |  |  |  |  | 294 | 224 | 56.76% |
| FM1046<br>(no<br>Auxin) | KLC-1 |  |  | yes |  |  |  |  |  | 538 | 3 | 99.84% |
| FM1046<br>(with<br>Auxin) | KLC-1 |  |  | yes |  |  |  |  |  | 393 | 8 | 97.78% |
| FM1165<br>(no<br>Auxin) | KLC-1,2 |  |  | yes |  |  |  |  |  | 450 | 1 | 99.78% |
| FM1165<br>(with<br>Auxin) | KLC-1,2 |  |  | yes |  |  |  |  |  | 171 | 121 | 58.56% |

**Table S2:** *C. elegans* strain list

| Strain # | Genotype | Figure |
| --- | --- | --- |
| FM691 | *cox-4(zu476[cox-4::eGFP::3xFLAG]) I; wjls76[Cn_unc-119(+); pie-1p::mKate2::tba-2]* | 1A |
| FM776 | *cox-4(zu476[cox-4::eGFP::3xFLAG]) I; unc-116(gk5722[loxP + myo-2p::GFP::unc-54 3' UTR + rps-27p::neoR::unc-54 3' UTR + loxP])III; duls1[unc-116::GFP integrated array]; wjls76[Cn_unc-119(+); pie-1p::mKate2::tba-2]* | 1B,1C |
| FM638 | *unc-119(ed3) III; ojls23 wjls76[Cn_unc-119(+); pie-1p::mKate2::tba-2]* | 1D |
| FM740 | *unc-116(f130) unc-36 (e251)III; ojls23[GFP::SP12; unc-119(+)]; wjls76[Cn_unc-119(+); pie-1p::mKate2::tba-2]* | 1E,1F |
| FM1165/<br>PHX8305 | *cpls103[sun-1p::TIR1::F2A::mTagBFP2::degron-NLS::tbb-2 3'UTR]II; KLC-1(syb7801[klc-1::AID::GFP])IV; KLC-2(syb8305[klc-2::AID::GFP])V; wjls76[Cn_unc-119(+); pie-1p::mKate2::tba-2]* | 1G,1H |
| FM1080/<br>PHX7873 | *kca-1(syb7873[kca-1::AID::GFP])I; cpls103[sun-1p::TIR1::F2A::mTagBFP2::degron-NLS::tbb-2 3'UTR]II; wjls76[Cn_unc-119(+); pie-1p::mKate2::tba-2]* | 1I,1J |
| FM955 | *Itls44[pAA173; pie-1p-mCh::PH(PLC1delta1) + unc-119(+)]V; vit-2(crg9070[vit-2::gfp])X; wjls76[Cn_unc-119(+); pie-1p::mKate2::tba-2]* | 2A |
| FM953,954 | *unc-116(f130) unc-36 (e251)III; vit-2(crg9070[vit-2::gfp])X; wjls76[Cn_unc-119(+); pie-1p::mKate2::tba-2]* | 2B |
| FM1089 | *klc-1(syb7801[klc-1::AID::GFP]); Itls44[pAA173; pie-1p-mCh::PH(PLC1delta1) + unc-119(+)]V; vit-2(syb5705[vit-2::HALO])X; wjls76[Cn_unc-119(+); pie-1p::mKate2::tba-2]* | 2E,2F, S1 |
| FM1094 | *kca-1(syb7873[kca-1::AID::GFP])I; him-8(e1489)IV; Itls44[pAA173; pie-1p-mCh::PH(PLC1delta1) + unc-119(+)]V; vit-2(syb5705[vit-2::HALO])X; wjls76[Cn_unc-119(+); pie-1p::mKate2::tba-2]* | 2C,2D,2H-K |
| FM1189 | *duSi16[pFM1973(TOMM-20::GFP-GLO::SSPBnanoGLO)]II; Itls44pAA173; [pie-1p-mCh::PH(PLC1delta1) + unc-119(+)]V; wjls76[Cn_unc-119(+); pie-1p::mKate2::tba-2]* | 3C |
| FM1185 | *duSi16[pFM1973(TOMM-20::GFP-GLO::SSPBnanoGLO)]II; unc-116(gk5722[loxP + myo-2p::GFP::unc-54 3' UTR + rps-27p::neoR::unc-54 3' UTR + loxP])III; duls1[unc-116::GFP integrated array]; wjls76[Cn_unc-119(+); pie-1p::mKate2::tba-2]* | 3D, S2 |
| FM1131 | *unc-116(gk5722[loxP + myo-2p::GFP::unc-54 3' UTR + rps-27p::neoR::unc-54 3' UTR + loxP])III; duSi14[pFM1953(K420::mKateGLO::iLID)]IV;* | 3E, S2, 4C |
|  | *Itls44pAA173; [pie-1p-mCh::PH(PLC1delta1) + unc-119(+)]V; duls1[unc-116::GFP integrated array]; wjls76[Cn_unc-119(+); pie-1p::mKate2::tba-2]* |  |
| FM1141 | *duSi16[pFM1973(TOMM-20::GFP-GLO::SSPBnanoGLO)]II; unc-116(gk5722[loxP + myo-2p::GFP::unc-54 3' UTR + rps-27p::neoR::unc-54 3' UTR + loxP])III; duSi14[pFM1953(K420::mKateGLO::iLID)]IV; Itls44pAA173; [pie-1p-mCh::PH(PLC1delta1) + unc-119(+)]V; duls1[unc-116::GFP integrated array]; wjls76[Cn_unc-119(+); pie-1p::mKate2::tba-2]* | 3F-3H, S3 |
| FM1137 | *duSi17[pFM1964(LMP-1::GFP-GLO::SSPBnanoGLO)]II; wjls76[Cn_unc-119(+); pie-1p::mKate2::tba-2]* | 4A |
| FM1139 | *duSi17[pFM1964(LMP-1::GFP-GLO::SSPBnanoGLO)]II; unc-116(gk5722[loxP + myo-2p::GFP::unc-54 3' UTR + rps-27p::neoR::unc-54 3' UTR + loxP])III; duls1[unc-116::GFP integrated array]; wjls76[Cn_unc-119(+); pie-1p::mKate2::tba-2]* | 4B |
| FM1156 | *duSi17[pFM1964(LMP-1::GFP-GLO::SSPBnanoGLO)]II; unc-116(gk5722[loxP + myo-2p::GFP::unc-54 3' UTR + rps-27p::neoR::unc-54 3' UTR + loxP])III; duSi14[pFM1953(K420::mKateGLO::iLID)]IV; Itls44pAA173; [pie-1p-mCh::PH(PLC1delta1) + unc-119(+)]V; duls1[unc-116::GFP integrated array]; wjls76[Cn_unc-119(+); pie-1p::mKate2::tba-2]* | 4D-F, S4 |
| FM1152 | *duSi31[pFM1968 (TMEM-147::GFP::SSPB(nanoGLO)]II; unc-116(gk5722[loxP + myo-2p::GFP::unc-54 3' UTR + rps-27p::neoR::unc-54 3' UTR + loxP])III; duSi14[pFM1953(K420::mKateGLO::iLID)]IV; Itls44pAA173; [pie-1p-mCh::PH(PLC1delta1) + unc-119(+)]V; duls1[unc-116::GFP integrated array]; wjls76[Cn_unc-119(+); pie-1p::mKate2::tba-2]* | 4H-J, S4 |
| FM1159 | *duSi30[pFM2034::mex5p::ebp-2::GFP(GLO)::SSPB(nano)::tbb2 3'UTR]II; unc-116(gk5722[loxP + myo-2p::GFP::unc-54 3' UTR + rps-27p::neoR::unc-54 3' UTR + loxP])III; duSi14[pFM1953(K420::mKateGLO::iLID)]IV; Itls44pAA173; [pie-1p-mCh::PH(PLC1delta1) + unc-119(+)]V; duls1[unc-116::GFP integrated array]; wjls76[Cn_unc-119(+); pie-1p::mKate2::tba-2]* | 4K-M, S4 |
| FM1154 | *egxSi126 [mex-5p::hsp-3(aa1-19)::halotag::HDEL::pie-1 3'UTR + unc-119(+)]I; Itls44pAA173; [pie-1p-mCh::PH(PLC1delta1) + unc-119(+)]V; vit-2(crg9070[vit-2::gfp])X; wjls76[Cn_unc-119(+); pie-1p::mKate2::tba-2];* | 5A |
| FM1055 | *egxSi126 [mex-5p::hsp-3(aa1-19)::halotag::HDEL::pie-1 3'UTR + unc-119(+)] I; duSi12[pFM1944(K420::mKateGLO::Khinge tail)]II; him-8(e1489)IV; Itls44pAA173; [pie-1p-mCh::PH(PLC1delta1) + unc-119(+)]V; vit-2(crg9070[vit-2::gfp]) X; wjls76[Cn_unc-119(+); pie-1p::mKate2::tba-2]* | 5B, 5D |
| FM1192 | *egxSi126 [mex-5p::hsp-3(aa1-19)::halotag::HDEL::pie-1 3'UTR + unc-119(+)] I; duSi12[pFM1944(K420::mKateGLO::Khinge tail)]II; unc-116(gk5722[loxP + myo-2p::GFP::unc-54 3' UTR + rps-27p::neoR::unc-54 3' UTR + loxP])III; vit-2(crg9070[vit-2::gfp]) X; duls1[unc-116::GFP integrated array]; wjls76[Cn_unc-119(+); pie-1p::mKate2::tba-2]* | 5C, |
| FM1193 | *cox-4(zu476[cox-4::eGFP::3xFLAG])I; duSi12[pFM1944(K420::mKateGLO::Khinge tail)]II; him-8(e1489)IV; Itls44pAA173; [pie-1p-mCh::PH(PLC1delta1) + unc-119(+)]V; wjls76[Cn_unc-119(+); pie-1p::mKate2::tba-2]* | 5E |
| FM1222 | *cox-4(zu476[cox-4::eGFP::3xFLAG])I; duSi12[pFM1944(K420::mKateGLO::Khinge tail)]II; unc-116(gk5722[loxP + myo-2p::GFP::unc-54 3' UTR + rps-27p::neoR::unc-54 3' UTR + loxP])III; Itls44pAA173; [pie-1p-mCh::PH(PLC1delta1) + unc-119(+)]V; duls1[unc-116::GFP integrated array]; wjls76[Cn_unc-119(+); pie-1p::mKate2::tba-2]* | 5F |
| FM1096 | *kca-1(syb7873) I; cpls103[sun-1p::TIR1::F2A::mTagBFP2::degron-NLS::tbb-2 3'UTR]II; sdhc-1(jbm1 [sdhc-1::mCherry]) III; wjls76[Cn_unc-119(+); pie-1p::mKate2::tba-2]* | S1 |
| FM1092 | *1p::TIR1::F2A::mTagBFP2::degron-NLS::tbb-2 3'UTR]II; sdhc-1(jbm1 [sdhc-1::mCherry]) III; klc-1(syb7801) IV; wjls76[Cn_unc-119(+); pie-1p::mKate2::tba-2]* | S1 |
| FM1210 | *duSi16[pFM1973(TOMM-20::GFP-GLO::SSPBnanoGLO)]II; duSi14[pFM1953(K420::mKateGLO::iLID)]IV; Itls44pAA173; [pie-1p-mCh::PH(PLC1delta1) + unc-119(+)]V; vit-2(syb5705[halo-tag])X; wjls76[Cn_unc-119(+); pie-1p::mKate2::tba-2]* | S3 |
| FM1214 | *duSi16[pFM1973(TOMM-20::GFP-GLO::SSPBnanoGLO)]II;*<br>*unc-116(gk5722[loxP + myo-2p::GFP::unc-54 3' UTR + rps-27p::neoR::unc-54 3' UTR + loxP])III;*<br>*duSi14[pFM1953(K420::mKateGLO::iLID)]IV; ItIs44pAA173;*<br>*[pie-1p-mCh::PH(PLC1delta1) + unc-119(+)]V; vit-2(syb5705[halo-tag])X; duls1[unc-116::GFP integrated array from FM75]; wjls76[Cn_unc-119(+); pie-1p::mKate2::tba-2]* | S3 |

## References

1. Hassold T, Hunt P. To err (meiotically) is human: the genesis of human aneuploidy. Nat Rev Genet. 2001;2(4):280–91. doi: 10.1038/35066065. PubMed PMID: 11283700.

2. Maddox AS, Azoury J, Dumont J. Polar body cytokinesis. Cytoskeleton (Hoboken). 2012;69(11):855–68. Epub 20121001. doi: 10.1002/cm.21064. PubMed PMID: 22927361.

3. Almonacid M, Terret M, Verlhac MH. Actin-based spindle positioning: new insights from female gametes. J Cell Sci. 2014;127(Pt 3):477–83. Epub 20140110. doi: 10.1242/jcs.142711. PubMed PMID: 24413163.

4. Yang HY, McNally K, McNally FJ. MEI-1/katanin is required for translocation of the meiosis I spindle to the oocyte cortex in C elegans. Dev Biol. 2003;260(1):245–59. Epub 2003/07/30. doi: 10.1016/s0012-1606(03)00216-1. PubMed PMID: 12885567.

5. Yang HY, Mains PE, McNally FJ. Kinesin-1 mediates translocation of the meiotic spindle to the oocyte cortex through KCA-1, a novel cargo adapter. J Cell Biol. 2005;169(3):447–57. doi: 10.1083/jcb.200411132. PubMed PMID: 15883196; PMCID: PMC2171918.

6. Ellefson ML, McNally FJ. CDK-1 inhibits meiotic spindle shortening and dynein- dependent spindle rotation in C. elegans. J Cell Biol. 2011;193(7):1229–44. Epub 2011/06/22. doi: 10.1083/jcb.201104008. PubMed PMID: 21690306; PMCID: PMC3216336.

7. Ellefson ML, McNally FJ. Kinesin-1 and cytoplasmic dynein act sequentially to move the meiotic spindle to the oocyte cortex in Caenorhabditis elegans. Mol Biol Cell. 2009;20(11):2722–30. Epub 2009/04/10. doi: 10.1091/mbc.E08-12-1253. PubMed PMID: 19357192; PMCID: PMC2688551.

8. van der Voet M, Berends CW, Perreault A, Nguyen-Ngoc T, Gonczy P, Vidal M, Boxem M, van den Heuvel S. NuMA-related LIN-5, ASPM-1, calmodulin and dynein promote meiotic spindle rotation independently of cortical LIN-5/GPR/Galpha. Nat Cell Biol. 2009;11(3):269–77. Epub 2009/02/17. doi: 10.1038/ncb1834. PubMed PMID: 19219036.

9. McCarter J, Bartlett B, Dang T, Schedl T. On the control of oocyte meiotic maturation and ovulation in Caenorhabditis elegans. Dev Biol. 1999;205(1):111–28. Epub 1999/01/12. doi: 10.1006/dbio.1998.9109. PubMed PMID: 9882501.

10. Vargas E, McNally KP, Cortes DB, Panzica MT, Danlasky BM, Li Q, Maddox AS, McNally FJ. Spherical spindle shape promotes perpendicular cortical orientation by preventing isometric cortical pulling on both spindle poles during C. elegans female meiosis. Development. 2019;146(20). Epub 2019/10/03. doi: 10.1242/dev.178863. PubMed PMID: 31575646; PMCID: PMC6826043.

11. Crowder ME, Flynn JR, McNally KP, Cortes DB, Price KL, Kuehnert PA, Panzica MT, Andaya A, Leary JA, McNally FJ. Dynactin-dependent cortical dynein and spherical spindle shape correlate temporally with meiotic spindle rotation in Caenorhabditis elegans. Mol Biol Cell. 2015;26(17):3030–46. Epub 20150701. doi: 10.1091/mbc.E15-05-0290. PubMed PMID: 26133383; PMCID: PMC4551317.

12. Panzica MT, Marin HC, Reymann AC, McNally FJ. F-actin prevents interaction between sperm DNA and the oocyte meiotic spindle in. J Cell Biol. 2017;216(8):2273–82. Epub 20170621. doi: 10.1083/jcb.201702020. PubMed PMID: 28637747; PMCID: PMC5551714.

13. McNally KL, Martin JL, Ellefson M, McNally FJ. Kinesin-dependent transport results in polarized migration of the nucleus in oocytes and inward movement of yolk granules in meiotic embryos. Dev Biol. 2010;339(1):126–40. Epub 2009/12/29. doi: 10.1016/j.ydbio.2009.12.021. PubMed PMID: 20036653; PMCID: PMC2823969.

14. Beath EA, Bailey C, Mahantesh Magadam M, Qiu S, McNally KL, McNally FJ. Katanin, kinesin-13, and ataxin-2 inhibit premature interaction between maternal and paternal genomes in. Elife. 2024;13. Epub 20240730. doi: 10.7554/eLife.97812. PubMed PMID: 39078879; PMCID: PMC11288632.

15. Cross JA, Dodding MP. Motor-cargo adaptors at the organelle-cytoskeleton interface. Curr Opin Cell Biol. 2019;59:16–23. Epub 20190402. doi: 10.1016/j.ceb.2019.02.010. PubMed PMID: 30952037.

16. Wu Y, Ding C, Sharif B, Weinreb A, Swaim G, Hao H, Yogev S, Watanabe S, Hammarlund M. Polarized localization of kinesin-1 and RIC-7 drives axonal mitochondria anterograde transport. J Cell Biol. 2024;223(5). Epub 20240312. doi: 10.1083/jcb.202305105. PubMed PMID: 38470363; PMCID: PMC10932739.

17. Zhao Y, Song E, Wang W, Hsieh CH, Wang X, Feng W, Shen K. Metaxins are core components of mitochondrial transport adaptor complexes. Nat Commun. 2021;12(1):83. Epub 20210104. doi: 10.1038/s41467-020-20346-2. PubMed PMID: 33397950; PMCID: PMC7782850.

18. Meyerzon M, Fridolfsson HN, Ly N, McNally FJ, Starr DA. UNC-83 is a nuclear-specific cargo adaptor for kinesin-1-mediated nuclear migration. Development. 2009;136(16):2725–33. Epub 20090715. doi: 10.1242/dev.038596. PubMed PMID: 19605495; PMCID: PMC2730402.

19. McNally KL, Fabritius AS, Ellefson ML, Flynn JR, Milan JA, McNally FJ. Kinesin-1 prevents capture of the oocyte meiotic spindle by the sperm aster. Dev Cell. 2012;22(4):788–98. Epub 20120329. doi: 10.1016/j.devcel.2012.01.010. PubMed PMID: 22465668; PMCID: PMC3606814.

20. Au V, Li-Leger E, Raymant G, Flibotte S, Chen G, Martin K, Fernando L, Doell C, Rosell FI, Wang S, Edgley ML, Rougvie AE, Hutter H, Moerman DG. CRISPR/Cas9 Methodology for the Generation of Knockout Deletions in. G3 (Bethesda). 2019;9(1):135-44. Epub 20190109. doi: 10.1534/g3.118.200778. PubMed PMID: 30420468; PMCID: PMC6325907.

21. Grant B, Hirsh D. Receptor-mediated endocytosis in the Caenorhabditis elegans oocyte. Mol Biol Cell. 1999;10(12):4311–26. doi: 10.1091/mbc.10.12.4311. PubMed PMID: 10588660; PMCID: PMC25760.

22. Poteryaev D, Fares H, Bowerman B, Spang A. Caenorhabditis elegans SAND-1 is essential for RAB-7 function in endosomal traffic. EMBO J. 2007;26(2):301–12. Epub 20070104. doi: 10.1038/sj.emboj.7601498. PubMed PMID: 17203072; PMCID: PMC1783445.

23. Tan Z, Yue Y, Leprevost F, Haynes S, Basrur V, Nesvizhskii AI, Verhey KJ, Cianfrocco MA. Autoinhibited kinesin-1 adopts a hierarchical folding pattern. Elife. 2023;12. Epub 20231101. doi: 10.7554/eLife.86776. PubMed PMID: 37910016; PMCID: PMC10619981.

24. Nijenhuis W, van Grinsven MMP, Kapitein LC. An optimized toolbox for the optogenetic control of intracellular transport. J Cell Biol. 2020;219(4). doi: 10.1083/jcb.201907149. PubMed PMID: 32328628; PMCID: PMC7147098.

25. Shcherbo D, Merzlyak EM, Chepurnykh TV, Fradkov AF, Ermakova GV, Solovieva EA, Lukyanov KA, Bogdanova EA, Zaraisky AG, Lukyanov S, Chudakov DM. Bright far-red fluorescent protein for whole-body imaging. Nat Methods. 2007;4(9):741–6. Epub 20070826. doi: 10.1038/nmeth1083. PubMed PMID: 17721542.

26. Perry AJ, Rimmer KA, Mertens HD, Waller RF, Mulhern TD, Lithgow T, Gooley PR. Structure, topology and function of the translocase of the outer membrane of mitochondria. Plant Physiol Biochem. 2008;46(3):265–74. Epub 20080103. doi: 10.1016/j.plaphy.2007.12.012. PubMed PMID: 18272380.

27. Gong T, McNally KL, Konanoor S, Peraza A, Bailey C, Redemann S, McNally FJ. Roles of Tubulin Concentration during Prometaphase and Ran-GTP during Anaphase of. Life Sci Alliance. 2024;7(9). Epub 20240703. doi: 10.26508/lsa.202402884. PubMed PMID: 38960623; PMCID: PMC11222656.

28. Lu W, Winding M, Lakonishok M, Wildonger J, Gelfand VI. Microtubule-microtubule sliding by kinesin-1 is essential for normal cytoplasmic streaming in Drosophila oocytes. Proc Natl Acad Sci U S A. 2016;113(34):E4995-5004. Epub 20160810. doi: 10.1073/pnas.1522424113. PubMed PMID: 27512034; PMCID: PMC5003289.

29. Duan X, Li Y, Yi K, Guo F, Wang H, Wu PH, Yang J, Mair DB, Morales EA, Kalab P, Wirtz D, Sun SX, Li R. Dynamic organelle distribution initiates actin-based spindle migration in mouse oocytes. Nat Commun. 2020;11(1):277. Epub 20200114. doi: 10.1038/s41467-019-14068-3. PubMed PMID: 31937754; PMCID: PMC6959240.

30. Cheng S, Altmeppen G, So C, Welp LM, Penir S, Ruhwedel T, Menelaou K, Harasimov K, Stützer A, Blayney M, Elder K, Möbius W, Urlaub H, Schuh M. Mammalian oocytes store mRNAs in a mitochondria-associated membraneless compartment. Science. 2022;378(6617):eabq4835. Epub 20221021. doi: 10.1126/science.abq4835. PubMed PMID: 36264786.

31. Beams HW, Kessel RG, Shih CY, Tung HN. Scanning electron microscopy studies on blastodisc formation in the zebrafish, Brachidanio rerio. Journal of Morphology. 1985;184:41–9. doi: 10.1002/jmor.1051840105.

32. Fernández J, Valladares M, Fuentes R, Ubilla A. Reorganization of cytoplasm in the zebrafish oocyte and egg during early steps of ooplasmic segregation. Dev Dyn. 2006;235(3):656–71. doi: 10.1002/dvdy.20682. PubMed PMID: 16425221.

33. Fernandez J, Olea N, Ubilla A, Cantillana V. Formation of polar cytoplasmic domains (teloplasms) in the leech egg is a three-step segregation process. Int J Dev Biol. 1998;42(2):149–62. PubMed PMID: 9551860.

34. Olson SK, Greenan G, Desai A, Müller-Reichert T, Oegema K. Hierarchical assembly of the eggshell and permeability barrier in C. elegans. J Cell Biol. 2012;198(4):731–48. doi: 10.1083/jcb.201206008. PubMed PMID: 22908315; PMCID: PMC3514041.

35. Nashchekin D, Fernandes AR, St Johnston D. Patronin/Shot Cortical Foci Assemble the Noncentrosomal Microtubule Array that Specifies the Drosophila Anterior-Posterior Axis. Dev Cell. 2016;38(1):61–72. doi: 10.1016/j.devcel.2016.06.010. PubMed PMID: 27404359; PMCID: PMC4943857.

36. Khuc Trong P, Doerflinger H, Dunkel J, St Johnston D, Goldstein RE. Cortical microtubule nucleation can organise the cytoskeleton of Drosophila oocytes to define the anteroposterior axis. Elife. 2015;4. Epub 20150925. doi: 10.7554/eLife.06088. PubMed PMID: 26406117; PMCID: PMC4580948.

37. Pfeiffer DC, Gard DL. Microtubules in Xenopus oocytes are oriented with their minus- ends towards the cortex. Cell Motil Cytoskeleton. 1999;44(1):34–43. doi: 10.1002/(SICI)1097-0169(199909)44:1&lt;34::AID-CM3>3.0.CO;2-6. PubMed PMID: 10470017.

38. Wang S, Wu D, Quintin S, Green RA, Cheerambathur DK, Ochoa SD, Desai A, Oegema K. NOCA-1 functions with γ-tubulin and in parallel to Patronin to assemble non-centrosomal microtubule arrays in C. elegans. Elife. 2015;4:e08649. Epub 20150915. doi: 10.7554/eLife.08649. PubMed PMID: 26371552; PMCID: PMC4608005.

39. Toya M, Kobayashi S, Kawasaki M, Shioi G, Kaneko M, Ishiuchi T, Misaki K, Meng W, Takeichi M. CAMSAP3 orients the apical-to-basal polarity of microtubule arrays in epithelial cells. Proc Natl Acad Sci U S A. 2016;113(2):332–7. Epub 20151229. doi: 10.1073/pnas.1520638113. PubMed PMID: 26715742; PMCID: PMC4720291.

40. Cha BJ, Serbus LR, Koppetsch BS, Theurkauf WE. Kinesin I-dependent cortical exclusion restricts pole plasm to the oocyte posterior. Nat Cell Biol. 2002;4(8):592–8. doi: 10.1038/ncb832. PubMed PMID: 12134163.

41. Sanchez AD, Feldman JL. Microtubule-organizing centers: from the centrosome to non- centrosomal sites. Curr Opin Cell Biol. 2017;44:93–101. Epub 20160922. doi: 10.1016/j.ceb.2016.09.003. PubMed PMID: 27666167; PMCID: PMC5362366.

42. Zhou K, Rolls MM, Hall DH, Malone CJ, Hanna-Rose W. A ZYG-12-dynein interaction at the nuclear envelope defines cytoskeletal architecture in the C. elegans gonad. J Cell Biol. 2009;186(2):229–41. doi: 10.1083/jcb.200902101. PubMed PMID: 19635841; PMCID: PMC2717649.

43. Bobinnec Y, Fukuda M, Nishida E. Identification and characterization of Caenorhabditis elegans gamma-tubulin in dividing cells and differentiated tissues. J Cell Sci. 2000;113 Pt 21:3747–59. doi: 10.1242/jcs.113.21.3747. PubMed PMID: 11034903.

44. McNally K, Audhya A, Oegema K, McNally FJ. Katanin controls mitotic and meiotic spindle length. J Cell Biol. 2006;175(6):881–91. Epub 2006/12/21. doi: 10.1083/jcb.200608117. PubMed PMID: 17178907; PMCID: PMC2064698.

45. Londoño-Vásquez D, Rodriguez-Lukey K, Behura SK, Balboula AZ. Microtubule organizing centers regulate spindle positioning in mouse oocytes. Dev Cell. 2022;57(2):197–211.e3. Epub 20220113. doi: 10.1016/j.devcel.2021.12.011. PubMed PMID: 35030327; PMCID: PMC8792338.

46. Xie B, Zhang L, Zhao H, Bai Q, Fan Y, Zhu X, Yu Y, Li R, Liang X, Sun QY, Li M, Qiao J. Poly(ADP-ribose) mediates asymmetric division of mouse oocyte. Cell Res. 2018;28(4):462–75. Epub 20180220. doi: 10.1038/s41422-018-0009-7. PubMed PMID: 29463901; PMCID: PMC5939045.

47. Fielmich LE, Schmidt R, Dickinson DJ, Goldstein B, Akhmanova A, van den Heuvel S. Optogenetic dissection of mitotic spindle positioning in vivo. Elife. 2018;7. Epub 20180815. doi: 10.7554/eLife.38198. PubMed PMID: 30109984; PMCID: PMC6214656.

48. Nonet ML. Efficient Transgenesis in. Genetics. 2020;215(4):903-21. Epub 20200608. doi: 10.1534/genetics.120.303388. PubMed PMID: 32513816; PMCID: PMC7404237.

49. Timmons L, Court DL, Fire A. Ingestion of bacterially expressed dsRNAs can produce specific and potent genetic interference in Caenorhabditis elegans. Gene. 2001;263(1-2):103–12. doi: 10.1016/s0378-1119(00)00579-5. PubMed PMID: 11223248.

50. Kirby C, Kusch M, Kemphues K. Mutations in the par genes of Caenorhabditis elegans affect cytoplasmic reorganization during the first cell cycle. Dev Biol. 1990;142(1):203–15. Epub 1990/11/01. doi: 10.1016/0012-1606(90)90164-e. PubMed PMID: 2227096.

51. Rivera Gomez K, Schvarzstein M. Immobilization of nematodes for live imaging using an agarose pad produced with a Vinyl Record. MicroPubl Biol. 2018;2018. Epub 20180809. doi: 10.17912/QG0J-VT85. PubMed PMID: 32550397; PMCID: PMC7282523.

